# Dbf4-Dependent Kinase Finetunes INO80 Function at Chromosome Replication Origins

**DOI:** 10.1101/2024.06.10.598178

**Authors:** Priyanka Bansal, Shibojyoti Lahiri, Chandni Kumar, Lorenzo Galanti, Erika Chacin, María Ángeles Ortíz-Bazán, Marisa Müller, Petra Vizjak, Tobias Straub, Felix Müller-Planitz, Andrés Aguilera, Belen Gómez-González, Boris Pfander, Axel Imhof, Christoph F. Kurat

**Affiliations:** Biomedical Center Munich (BMC), Division of Molecular Biology, Faculty of Medicine, Ludwig-Maximilians-Universität in Munich, Martinsried, Germany; Cell Biology, Dortmund Life Science center (DOLCE), TU Dortmund University, Faculty of Chemistry and Chemical Biology, Dortmund, Germany; Present address: DSB Repair Metabolism Laboratory, The Francis Crick Institute, London, UK; Centro Andaluz de Biología Molecular y Medicina Regenerativa-CABIMER, Universidad de Sevilla-CSIC, Seville, Spain; Institute of Physiological Chemistry, Faculty of Medicine Carl Gustav Carus, Technische Universität Dresden, Dresden, Germany; Bayer AG, Pharmaceuticals, Research and Development, Genomic Medicine, Aprather Weg 18a, 42113 Wuppertal - Aprath, Germany; Early Stage Bioprocess Development, Boehringer Ingelheim Pharma GmbH & Co. KG, Biberach an der Riss, Germany

## Abstract

The highly conserved Dbf4-Dependent Kinase (DDK) plays a pivotal role in the nucleus during S phase, where it directly phosphorylates the replicative helicase, the minichromosome maintenance (MCM) complex. This leads to the initiation of chromosome replication. However, aside from the MCM complex, few other targets have been identified to date, leaving DDK an understudied kinase.

Here, we describe a two-pronged mass spectrometry-based approach and define the nuclear DDK-dependent phosphoproteome, which consists of approximately 400 phosphorylation events. Within this network, we found that DDK directly phosphorylates the Arp8 subunit of the multi-subunit chromatin remodeler complex INO80. Arp8 phosphorylation stabilises INO80’s intramolecular complex integrity, which finetunes its nucleosome spacing activity at replication origins. This adjustment of origin chromatin architecture stimulates replication and is important for the response to replication stress. Our results represent a significant advance in our understanding of the molecular mechanisms underlying the regulation of replication origins.

## Introduction

The replication of nuclear genomic DNA represents one of the most fundamental processes that a cell is capable of undertaking. Any failure to replicate nuclear genomic DNA can have significant consequences for genome integrity, with the potential to result in DNA damage, which is a hallmark of cancer development. It is therefore of the utmost importance that replication is highly regulated and that repair mechanisms are able to act quickly and efficiently if things go out of control. Genomic DNA is wrapped around nucleosomes, the basic units of chromatin, which provides a means for the safe storage and packaging of free DNA (Kornberg et al., 1999). However, nucleosomes are relatively stable, which presents a challenge for DNA-templated processes such as replication and transcription. For example, the replication machinery requires the assistance of the histone chaperone FACT/Nhp6a to replicate through a chromatin template in a biochemically reconstituted system (Kurat et al., 2017; Safaric et al., 2022). It is notable that nucleosome spacing remodelers, such as INO80 or ISW1a, were also found to be essential for achieving *in vivo*-like replication rates, indicating that nucleosome spacing plays a pivotal role (Kurat et al., 2017). Indeed, recent work has demonstrated that optimal nucleosome spacing around replication origins, facilitated by the ATP-dependent remodelers INO80, ISW1a, ISW2 and Chd1, is crucial for efficient replication initiation (Chacin et al., 2023).

The process of DNA replication is a two-step process (Bell et al., 2016; Costa et al., 2022). In the G1 phase of the cell cycle, the Origin Recognition Complex (ORC) loads the inactive MCM complex onto origins of replication with the assistance of other loading factors and ATP. In budding yeast, origins are short DNA sequences (Liachko et al., 2013; Nieduszynski et al., 2006; Siow et al., 2012; Xu et al., 2006) within a nucleosome-free region (NFR), which are flanked by the aforementioned well-positioned nucleosomes on both sites (Berbenetz et al., 2010; Eaton et al., 2010; Rossi et al., 2021). It is noteworthy that, in addition to its canonical role as the MCM loader, ORC plays a second essential role prior to replication initiation in the G1 phase (Chacin et al., 2023). This involves acting as a boundary element and cooperating with INO80, ISW1a, ISW2 and Chd1 to generate the chromatin landscape around origins.

Subsequently, during the S phase, the MCMs are activated in order to initiate the replication process (Bell et al., 2016; Costa et al., 2022). The number of origins is proportional to the size of the genome. For instance, budding yeast employs approximately 300 core origins to replicate its 2 mega base genome (Liachko et al., 2013; Nieduszynski et al., 2006; Siow et al., 2012; Xu et al., 2006), whereas in human cells, it is estimated that approximately 30,000 origins are required, which correlates with the roughly 100 fold larger human genome (Akerman et al., 2020). Origins do not fire simultaneously; rather, they follow a temporal programme, or replication timing (RT), where origins are divided into early- or late-firing (Creager et al., 2015; Czajkowsky et al., 2008; Dimitrova et al., 1999; Fragkos et al., 2015; Hawkins et al., 2013; Leonard et al., 2013; Raghuraman et al., 1997; Saner et al., 2013). The maintenance of the RT programme is crucial for genome integrity. However, the underlying molecular mechanisms remain poorly understood.

In addition to various replication factors, two prominent cell cycle kinases, namely Cyclin-Dependent Kinase (CDK) and Dbf4-Dependent Kinase (DDK), are responsible for transforming the inactive MCM complex into the active helicase, which initiates replication (Bell et al., 2016; Costa et al., 2022). CDK is a well-studied kinase with a multitude of target proteins involved in a plethora of cellular processes (Enserink et al., 2010; Morgan, 1995; Ubersax et al., 2003). Distinct kinase-activating proteins, called cyclins, oscillate and accumulate during all major cell cycle stages, ensuring that CDK is active throughout the cell cycle (Bloom et al., 2007). In contrast, DDK’s kinase Cdc7 has only one activator, Dbf4, which accumulate in S phase, where it is essential for the initiation of replication (Bousset et al., 1998; Donaldson et al., 1998; Gillespie et al., 2022; Jiang et al., 1999; Sheu et al., 2006, 2010). In addition to DDK’s pivotal role in S phase, Dbf4 levels remain high until the metaphase-to-anaphase transition and DDK is involved in processes like meiotic recombination (Sasanuma et al., 2008; Wan et al., 2008), chromosome segregation (Argunhan et al., 2017; Challa et al., 2019; He et al., 2020; Matos et al., 2008), DNA double-strand break repair (Galanti et al., 2024; Princz et al., 2017) and the block to over-replication (Reusswig et al., 2016). Interestingly, DDK was found to interact with ORC at early-replicating origins in the G1 phase (Duncker et al., 2002; Fang et al., 2017; Pasero et al., 1999). The biological significance of this interaction, however, remains to be elucidated. In contrast to CDK, only a limited number of target proteins have been identified, which continues to make DDK a somewhat enigmatic kinase.

The objective of this study was to gain further insights into the biology of DDK by elucidating a global DDK-dependent phosphorylation network of nuclear processes. To this end, we developed a two-pronged mass spectrometry-based screening pipeline. This involved the inhibition of DDK function *in vivo* via two independent methods, the enrichment of nuclear fractions and the analysis of the nuclear phospho-proteomes. A total of approximately 400 putative DDK substrates were identified, with the majority of the target proteins being associated with chromatin-templated processes.

Among the identified DDK target sites we identified two phosphorylation sites within an unstructured region of the Arp8 subunit of INO80. As we have previously identified a critical function of the INO80 complex in setting up a defined chromatin structure at replication origins (Chacin et al., 2023), we wondered whether the DDK-dependent phosphorylation of Arp8 might regulate this function of INO80. Indeed, a mutant INO80 complex bearing a Arp8-phospho mutant subunit, were all phosphorylated serine were converted into alanine residues, showed a significant structural rearrangement of important modules of INO80, which in turn resulted in a reduction of INO80’s intrinsic capability to hydrolyse ATP. In accordance with our hypothesis, DDK-mediated phosphorylation of Arp8 is crucial for the correct positioning of nucleosomes at replication origins. Together with the biochemical findings, our *in vivo* data demonstrated that the loss of Arp8 phosphorylation results in a significant replication defect and a severe growth phenotype under replication stress conditions.

Our findings collectively indicate that DDK exerts a profound influence on chromatin architecture at replication origins, as well as on origin function. This influence is mediated by a “finetuning mechanism” that precisely positions nucleosomes through the action of a nucleosome spacing remodeler. We propose a model whereby this mechanism may contribute to the RT programme and the maintenance of genome integrity.

## Results

### A global network of DDK phosphorylation in the nucleus

The objective of this study was to expand our knowledge about DDK-dependent phosphorylation events and to identify phosphoproteins using a global proteomic approach, which included phosphopeptide enrichment in the presence and absence of DDK function *in vivo.* A comparable approach was recently published, in which DDK phosphoproteomes were identified in M-phase-arrested budding yeast cells (Galanti et al., 2024). Given that our primary interest was in chromatin dynamics and associated processes during replication, we reasoned that it would be most informative to analyse phosphoproteomes of nuclear fractions and specifically determine phosphoproteomes of samples in S phase.

We then considered potential methods for inhibiting DDK. DDK is essential in yeast, so we employed the use of *cdc7-4* (On et al., 2014), a strain bearing a temperature-sensitive allele of the *CDC7* gene. This strain allows us to maintain DDK function when cells are grown at a permissive temperature (25 °C), while eliminating it at a restrictive temperature (37 °C) (Figure 1A). It can be expected that DDK-dependent phosphorylation events will be reduced or even absent upon inhibition of the kinase by growing the strain at a restrictive temperature. However, it should be noted that yeast has an optimal growth temperature of 30 °C, so both 25 and 37 °C may result in an increase in false-positive rates caused by sub-optimal temperature.

**Figure 1:**
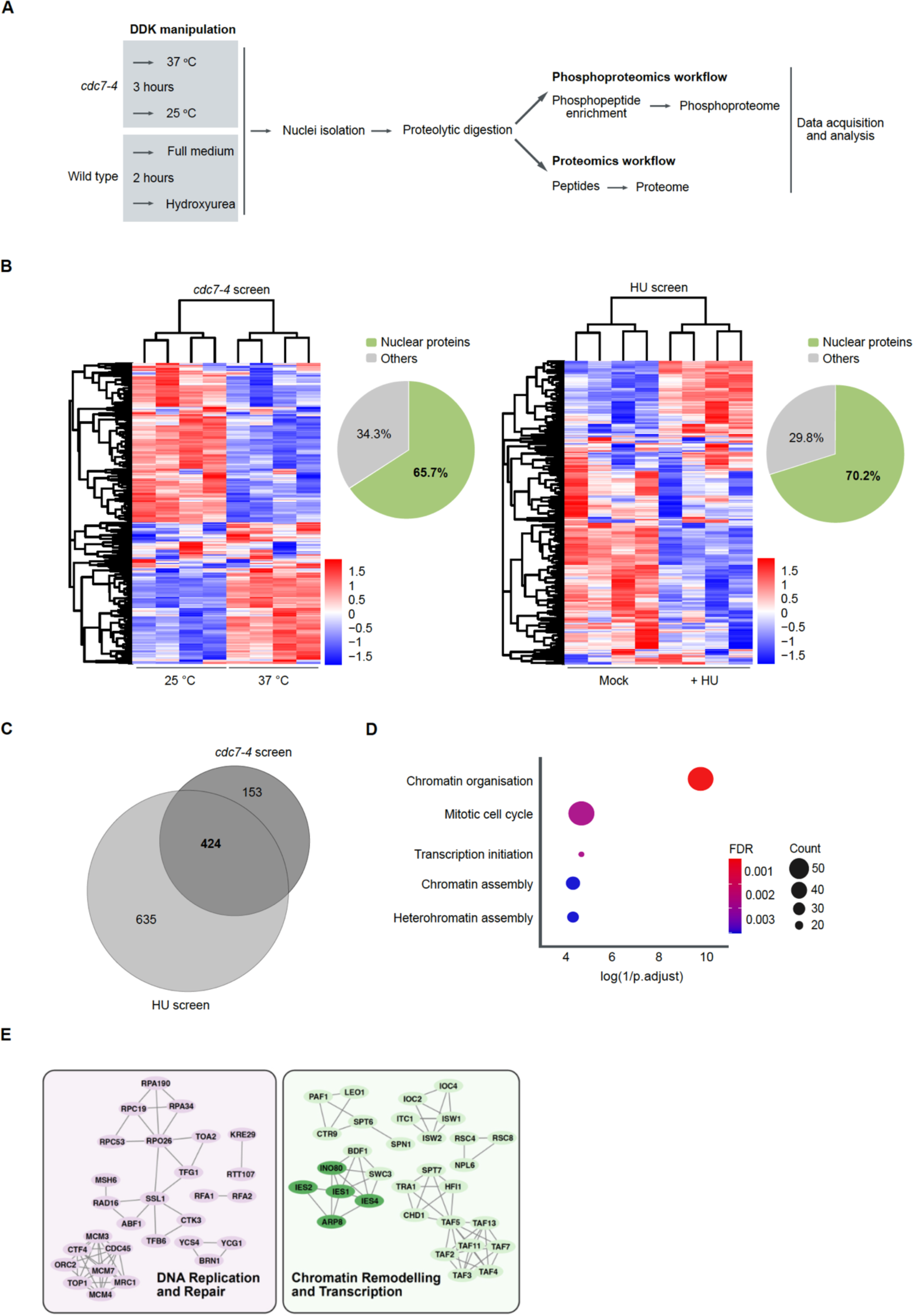
The nuclear DDK-dependent phospho-proteomic network. **A**) Schematic representation of the proteomic workflow used to enrich phospho-peptides to determine the phosphoproteome landscape upon perturbation of DDK activity. **B**) Heatmaps showing the clustering of top 500 variable phosphosites in the *cdc7-4* (left map) and Hydroxyurea screens (right map) respectively. Each column of the heatmap is a replicate of the corresponding condition (25 or 37°C; Plus (HU) or minus (Mock) HU). *Insets*: Pie charts showing nuclear enrichment of the phosphoproteins in the respective screens. The temperature screen shows an enrichment of 65.7%, whereas the HU screen shows 70.2% enrichment. N=4. **C**) Venn-diagram showing the overlap of enriched phosphosites in the DDK active fraction of the *cdc7-4* and HU screens. The comparative analysis was performed on all the down-regulated sites along with the exclusive sites. Sites were considered exclusive to a condition (e.g. permissive temperature or no HU treatment), when the perturbed condition (e.g. restrictive temperature or HU treatment) had the same sites detected in none or just 1 replicate. **D**) GO term analysis using the corresponding proteins of the 424 overlapping phosphosites. Plots for the top 5 most significant biological processes that are over-represented in the overlapping phosphosite dataset are shown here. Plot was generated using the R package ‘ggplot’. E) Shown are all bona fide nuclear proteins that show DDK-dependent phosphorylation in both screens and have at least one high confidence interactor. Grouping was done manually. Subunits of the INO80 complex are highlighted in dark green.

We therefore considered an independent method of inhibiting DDK. We reasoned that a comparison of both DDK-affected datasets would help to define a “high-confidence” DDK phosphorylation network. Another method of inhibiting DDK function, at least for its ability to phosphorylate the MCM complex, is the activation of the DNA damage checkpoint. An active checkpoint results in the binding of the checkpoint kinase Rad53 (mammalian Chk1) to DDK, phosphorylating it and thereby inhibiting its binding to the MCM complex (Abd Wahab et al., 2020; Greiwe et al., 2022; Lopez-Mosqueda et al., 2010; Zegerman et al., 2010). This prevents MCM phosphorylation and subsequent origin firing under replication stress conditions. Given our primary objective of identifying DDK targets that were phosphorylated under comparable conditions to the MCM complex, we experimentally triggered the checkpoint to influence DDK binding in a manner analogous to that observed for the MCM complex. Cells were treated with the replication drug hydroxyurea (HU), which decreases dNTP pools and thus causes replication stress. It is noteworthy that the *cdc7-4* mutants (Hereford et al., 1974), when incubated under restrictive temperature conditions or wild type cells treated with HU, exhibited a cell cycle arrest, whereas the control samples did not. This may also result in the indirect phosphorylation of other proteins.

As our primary focus was on the nuclear targets of DDK, we conducted a phosphoproteome analysis of nuclear extracts (Figure 1A) from *cdc7-4* cells under permissive or restrictive conditions (Figure 1B, left panel) and from cells that were either treated or not-treated with HU (Figure 1B, right panel). As anticipated, differential phosphorylation was observed under both conditions, consistent with the pivotal role of phosphorylation in regulating the cell cycle. In order to identify DDK-dependent phosphorylation sites, we focused our further analysis on the sites that were enriched in cells carrying an active DDK kinase in both screens. A Gene Ontology (GO) analysis of the 424 putative DDK targets (Figure 1D) revealed that chromatin organisation was one of the most significant processes regulated by DDK *in vivo*. A network analysis of all bona fide nuclear proteins that exhibited a clear increase in phosphorylation at permissive temperature (for the *cdc7-4* screen) or without HU (for the HU screen) and revealed several large complexes involved in DNA replication and transcription as putative targets of DDK (Figure 1E and S1). As we recently demonstrated that the INO80 complex plays a pivotal role in establishing a regular spacing of nucleosomes around replication origins (Chacin et al., 2023), we found it intriguing to observe that numerous complex components were phosphorylated when DDK was active (Figure 1E). However, the experimental setup does not permit the distinction between sites that are directly phosphorylated by DDK and sites that are phosphorylated by a downstream kinase or indirect consequences of the cell cycle arrest.

### The Arp8 subunit of INO80 is a bona fide DDK target

We next wanted to determine which of the INO80 subunits is a direct substrate of DDK. To achieve this, an *in vitro* kinase assay was conducted using the purified INO80 complex and DDK. INO80 is comprised of 15 subunits, with numerous serine and threonine residues present in the primary structures. Surprisingly, only one subunit, of approximately the same size as Arp8 (Figure 2A), undergoes a significant degree of phosphorylation in the presence of purified INO80 and DDK *in vitro* (Figure 2B, C and D). In accordance with this, two serine residues, namely serine 65 and 233, were identified as being significantly dependent on the presence of DDK in our phosphoproteome datasets (Figure 1 and 2A). The serine residues were mutated to alanine, and the resulting complex (INO80-AA) was purified (Figure 2A and B). The mutation of serine 65 and serine 233 of Arp8 did not result in a modified stoichiometric architecture of INO80-AA compared to INO80, as determined by SDS-PAGE followed by Coomassie staining (Figure 2B). However, it did result in strong reduction in the ability of DDK to phosphorylate Arp8 in the complex (Figure 2C and D). This indicates that (i) the Arp8 subunit of the INO80 complex is phosphorylated *in vitro* by DDK and that (ii) that serine 65 and serine 233 are the primary sites. The subsequent objective was to investigate the consequences of a loss of Arp8 phosphorylation by DDK. Having obtained purified complexes, the initial question was whether phosphorylation might influence INO80’s enzymatic activity to hydrolyse ATP. To address this question, an ATPase assay was employed. In this assay, ATP hydrolysis is coupled to the conversion of phosphoenolpyruvate to lactate. Lactate production consumes NADH, which is then measured photometrically and allows for the calculation of ATP hydrolysis rates (Figure 2E). Interestingly, we observed a significant reduction in the ATP hydrolysis rates with the INO80-AA mutant complex compared to INO80 wild type (Figure 2F). ATP hydrolysis was dependent on DNA and, in contrast to other remodelers, could not be stimulated by the addition of chromatin, in accordance with previous results with purified INO80 (Zhou et al., 2016).

**Figure 2:**
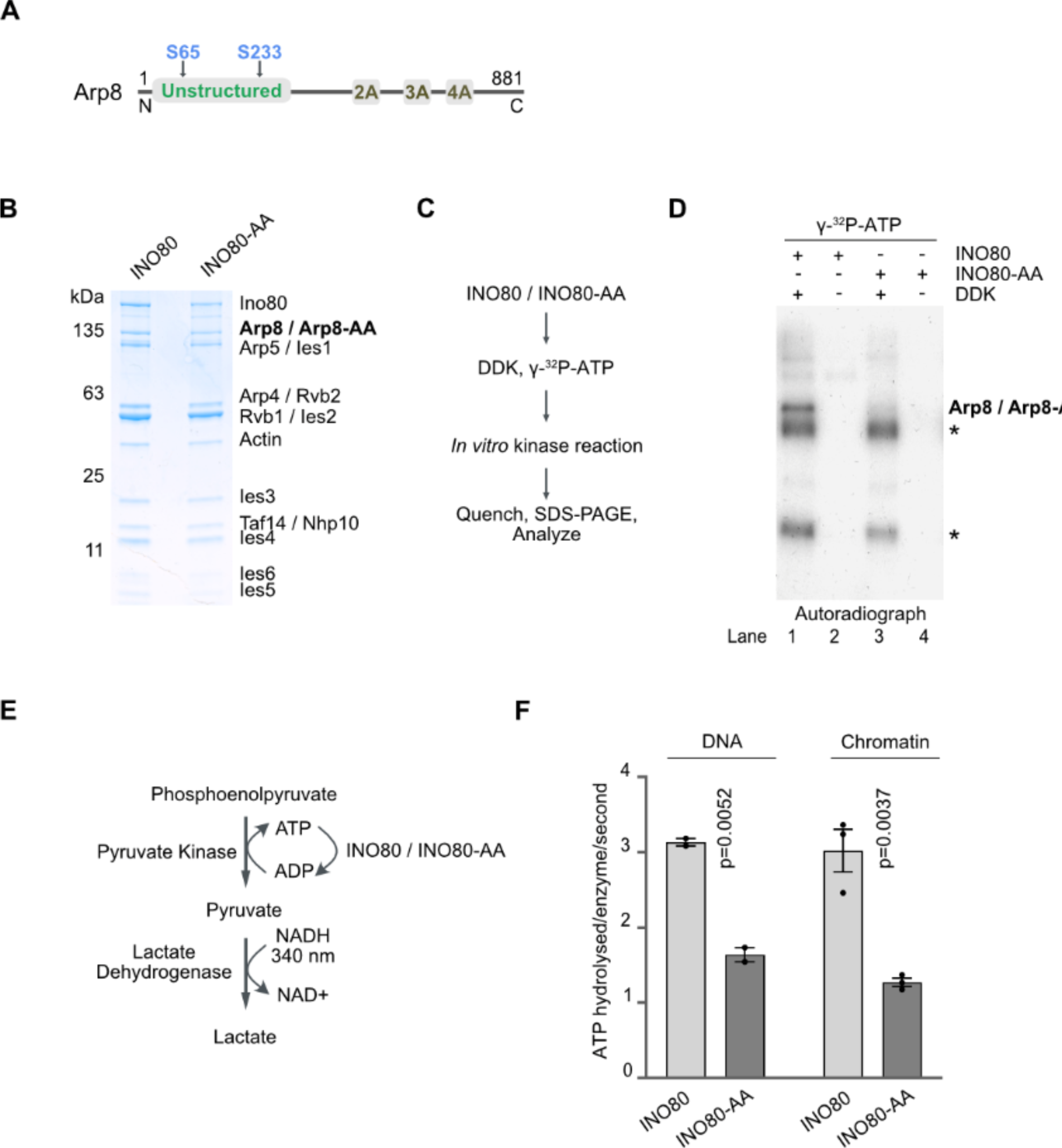
The Arp8 subunit of the INO80 complex is a bona-fide substrate of DDK. **A)** Domain organization of the Arp8 Subunit of the INO80 complex. Highlighted is the unstructured region and the two identified DDK-dependent phosphorylation sites serine 65 and serine 233. 2A, 3A and 4A represent non-actin folds. **B)** SDS-PAGE analysis of purified INO80 wild type and INO80-AA mutant complexes. **C)** Reaction scheme for the *in vitro* kinase reaction using purified DDK, INO80 and INO80-AA complexes. **D)** Incorporation of [^32^P]-γ-ATP into INO80 and INO80-AA by DDK was visualized by autoradiography after separation using SDS-PAGE. Asterisks show auto-phosphorylation of DDK. N=1. **E)** Outline of the assay to measure ATP hydrolysis rates. **F)** ATP hydrolysis rates by purified INO80 and INO80-AA in the presence of DNA (N=2) and chromatin (N=3) as substrates. ATP hydrolyzed per enzyme per second were obtained by measuring the oxidation of NADH, which is used to regenerate ADP to ATP. The graphs were plotted with standard mean and error and p-values were obtained by using two-tailed unpaired t-test calculations.

Our findings collectively indicate that DDK-dependent phosphorylation of the Arp8 subunit of INO80 does not impede the stoichiometric assembly of the complex, but is crucial for its intrinsic capacity to hydrolyse ATP.

### Arp8 phosphorylation is important for intramolecular integrity of INO80

INO80 is a ∼ 1 megadalton chromatin remodeler with at least 15 subunits and organised in three structural modules termed “N” (Nhp10/Ies1/Ies3/Ies5), “A” (Arp4/Arp8/Act1(Actin)/Taf14/Ies4) and “C” (Rvb1/Rvb2/Arp5/Ies2/Ies6) (Figure 3A) (Eustermann et al., 2018; Kunert et al., 2022; Tosi et al., 2013). The Ino80 subunit itself carries the ATPase motor activity and acts as a scaffold for the assembly of the three domains. The C module contains, among other proteins, the scaffolding AAA^+^-ATPases Rvb1 and Rvb2 and the ATPase motor domain of Ino80 (Ino80 insertion), which can interact with the nucleosome and pump extranucleosomal DNA into the nucleosome core particle as part of the sliding mechanism (Eustermann et al., 2018). The function of the N and A modules remains less well understood. It has been demonstrated that an interaction between the N module and the Ino80 scaffold is crucial for INO80’s spacing activity (Oberbeckmann et al., 2021; Zhou et al., 2018). Furthermore, the A module, which contains Arp8, has been described as being able to bind extranucleosomal DNA from the edge of the nucleosome (Brahma et al., 2018; Gerhold et al., 2012; Knoll et al., 2018; Kunert et al., 2022; Saravanan et al., 2012; Szerlong et al., 2008). This enables it to function as a DNA length sensor, regulating nucleosomal sliding by the motor ATPase Ino80. It is noteworthy that the N-terminal unstructured region of Arp8, encompassing serine 65 and 233 (Figure 2A), was demonstrated to be pivotal for this function (Brahma et al., 2018) and regulates the spacing activity of INO80 *in vivo* (Singh et al., 2021). This suggests a potential role for DDK-dependent phosphorylation of serine 65 and 233 not only in ATP hydrolysis by the Ino80 motor, but also in the regulation of nucleosomal sliding.

**Figure 3:**
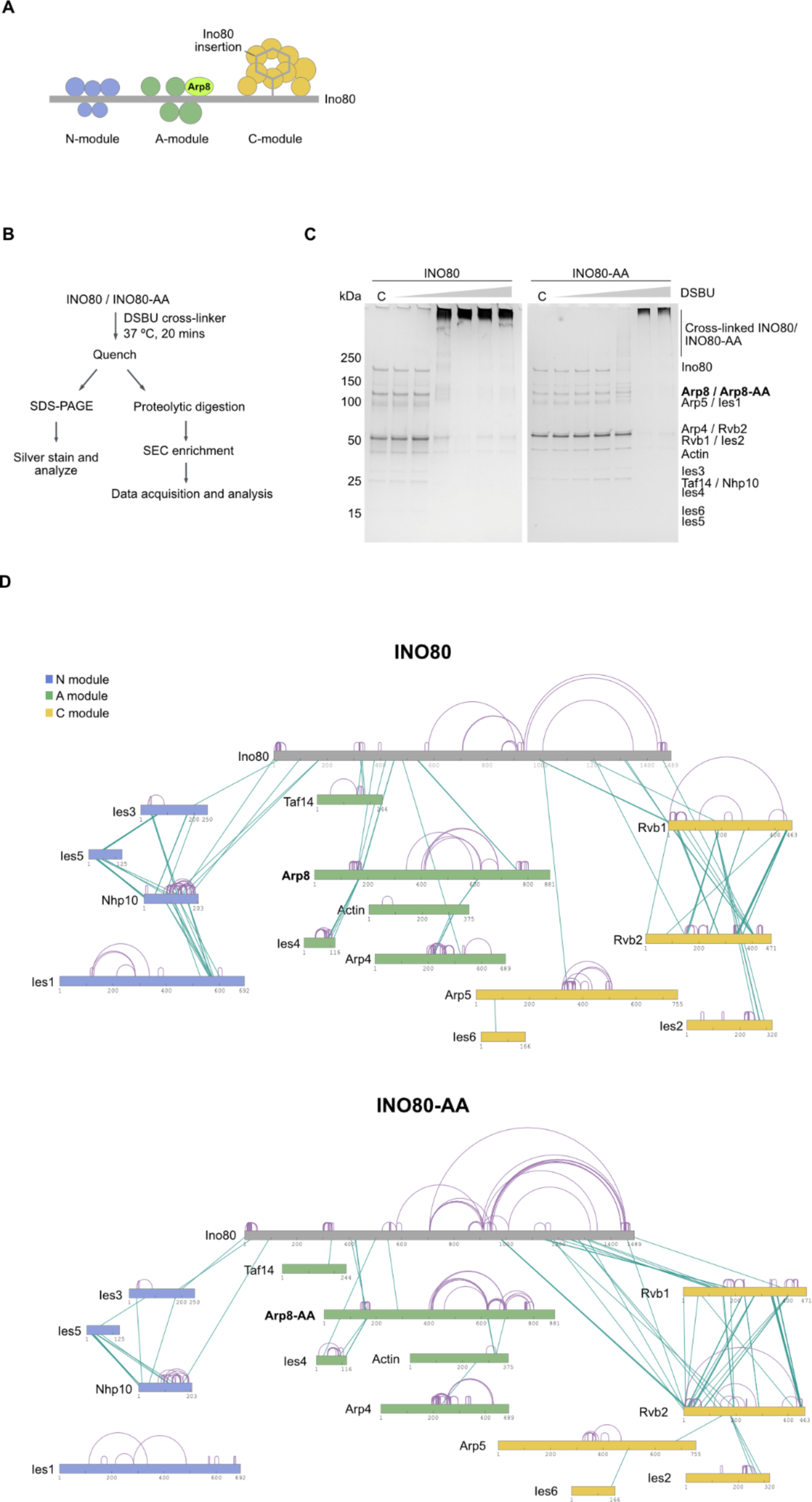
Subunit interaction topology maps of INO80 and INO80-AA complexes. **A)** Schematic depicting the organizational structure of the INO80 complex’s subunits and submodules. The Arp8 subunit is highlighted. The Submodules “N” (NHP10 module), “A” (ARP8 module) and “C” (Core module) are color coded in blue, green and yellow respectively. **B)** Outline of the cross-linking reactions using ureido-4,4’-dibutyric acid bis (hydroxysuccinimide) ester (DSBU) crosslinker with purified INO80 and INO80-AA complexes. **C)** SDS-PAGE and silver stain analysis of the DSBU crosslinker titration with INO80 and INO80-AA complexes. INO80 or INO80-AA complexes were tested with a range of 25 – 208 µM DSBU as represented by grey triangle on the top. C refers to control reaction without any crosslinker and kDa refers to the size of proteins in kilo-daltons. N=3. **D)** Crosslink network map of the INO80 and INO80-AA complexes. Different subunits and modules of INO80 are color coded in accordance with (A). Inter cross-links are represented in green and intra cross-links are represented in purple within the complex. N=3.

Given that no differences were observed in the stoichiometric composition of INO80 compared to INO80-AA, we hypothesised that Arp8 phosphorylation might be important for stabilising the sub-molecular composition of the complex, which, in turn, might be important for ATPase and nucleosomal spacing activity. However, this N-terminal part of INO80 was not amenable to resolution via cryo-electron microscopy (Eustermann et al., 2018). Consequently, we postulated that determining the subunit interaction topology of both complexes via cross-linking mass spectrometry would prove informative. While this approach offers a lower resolution compared to cryo-electron microscopy, it may provide insights into potential differences in sub-complex formation between the INO80 and INO80-AA mutant complexes.

As previously demonstrated (Tosi et al., 2013), the INO80 complex exhibits a distinctive topology of the individual modules. The objective was to investigate the potential for different topologies to form in the INO80-AA mutant. A similar protocol was therefore employed, as previously stated (Tosi et al., 2013) but using an MS2-cleavable DSBU crosslinker instead of an isotope-labeled DSS (Figure 3B). The optimal crosslinking concentration was determined by titration experiments. The two complexes exhibited different behaviors in the presence of the DSBU crosslinker, with the INO80 wild-type requiring lower concentrations of the crosslinker than the INO80-AA mutant complex (Figure 3C). This observation is indicative of structural differences between the two complexes.

We proceeded to apply mass spectrometry analyses of the crosslinked INO80 complexes. In the case of the INO80 wild-type complex, this led to the identification of approximately 300 intracrosslinks (within the same polypeptide) (Table S1) and 180 interlinks (between two different polypeptides) (Figure 3D, upper panel; Table S1). In accordance with previous work (Tosi et al., 2013), the three additional topological modules (“N”, “A” and “C”) were identified in addition to the Ino80 scaffold. The N module is crosslinked with the N-terminal part of Ino80, the A module with the middle part, and the C module with the C-terminal part. Notably, the number of intra- and intercrosslinks of the INO80-AA complex was found to be altered compared to INO80 (Figure 3D), in accordance with the SDS-PAGE results of the crosslinked complexes (Figure 3C). The crosslinks between the A module and Ino80 were reduced in the mutant complex, whereas they were increased between the C module and Ino80 scaffold. Most strikingly, the number of crosslinks between Nhp10 and Ino80 was significantly reduced, and between Ies1 and Nhp10 were virtually absent in the INO80-AA mutant complex compared to the wild-type (Figure 3D).

Collectively, our findings demonstrate that impaired phosphorylation of the Arp8 subunit of INO80 is correlated with a change in subunit topology. The interactions between the N module and the Ino80 scaffold were found to be most affected, leading us to hypothesise that phosphorylation of Arp8 might be crucial for INO80’s function in nucleosomal spacing.

### Arp8 phosphorylation is important for nucleosome spacing activity of INO80

A recent study has demonstrated that INO80 interacts with other chromatin remodellers, including ISW1a, ISW2 and Chd1, to establish regularly spaced nucleosomal arrays at replication origins (Chacin et al., 2023). A change in subunit topology was observed in the INO80-AA mutant complex, particularly between the N-terminal part of the Ino80 scaffold and the N module (Figure 3D). We therefore sought to ascertain whether DDK-dependent phosphorylation of Arp8 affects nucleosomal spacing at replication origins. To determine this, we employed a genome-scale *in vitro* assay, utilising purified histones and salt gradient dialysis (SGD) to assemble chromatin onto a library comprising approximately 300 individual budding yeast replication origins as previously described (Chacin et al., 2023) (Figure 4A). Subsequently, we conducted an assay of purified ORC in conjunction with INO80 wild-type and INO80-AA mutant complexes. A composite plot of the data indicates that our findings are consistent with those of our previous analyses (Chacin et al., 2023), which indicated that ORC and INO80 can generate a nucleosome-free region (NFR) at the ACS sites and assemble bi-directional and regularly spaced nucleosomes (Figure 4B). In contrast, while the INO80-AA mutant complex was able to form a nucleosome-free region (NFR) in a manner comparable to the wild-type complex, the positioning of the flanking nucleosomes (+ and – nucleosomes) was less pronounced (Figure 4B). Furthermore, the spacing patterns (internucleosomal linker length) downstream of the + and – nucleosomes exhibited differences. The INO80-AA mutant generated a wider linker length compared to the INO80 wild type. In addition to the general trends observed in the ACS-aligned composite plots, we also employed ACS-aligned heat maps to illustrate the variation among individual origins. The heat maps were sorted by the regularity of the NFR-array pattern for each condition. The majority of the origins (∼80%) exhibited the NFR-array pattern when ORC was combined with the INO80 wild-type complex (Figure 4C). The few origins that did not show this pattern may have different affinities for ORC binding and/or may require additional factors, which will require further investigation. In contrast, the INO80-AA mutant complex demonstrated that a significantly greater number of origins did not adopt the regular NFR-array pattern. Furthermore, even within the top-ranked origins in terms of regularity, the nucleosomal spacing appeared to be less defined compared to the wild type (Figure 4C).

**Figure 4:**
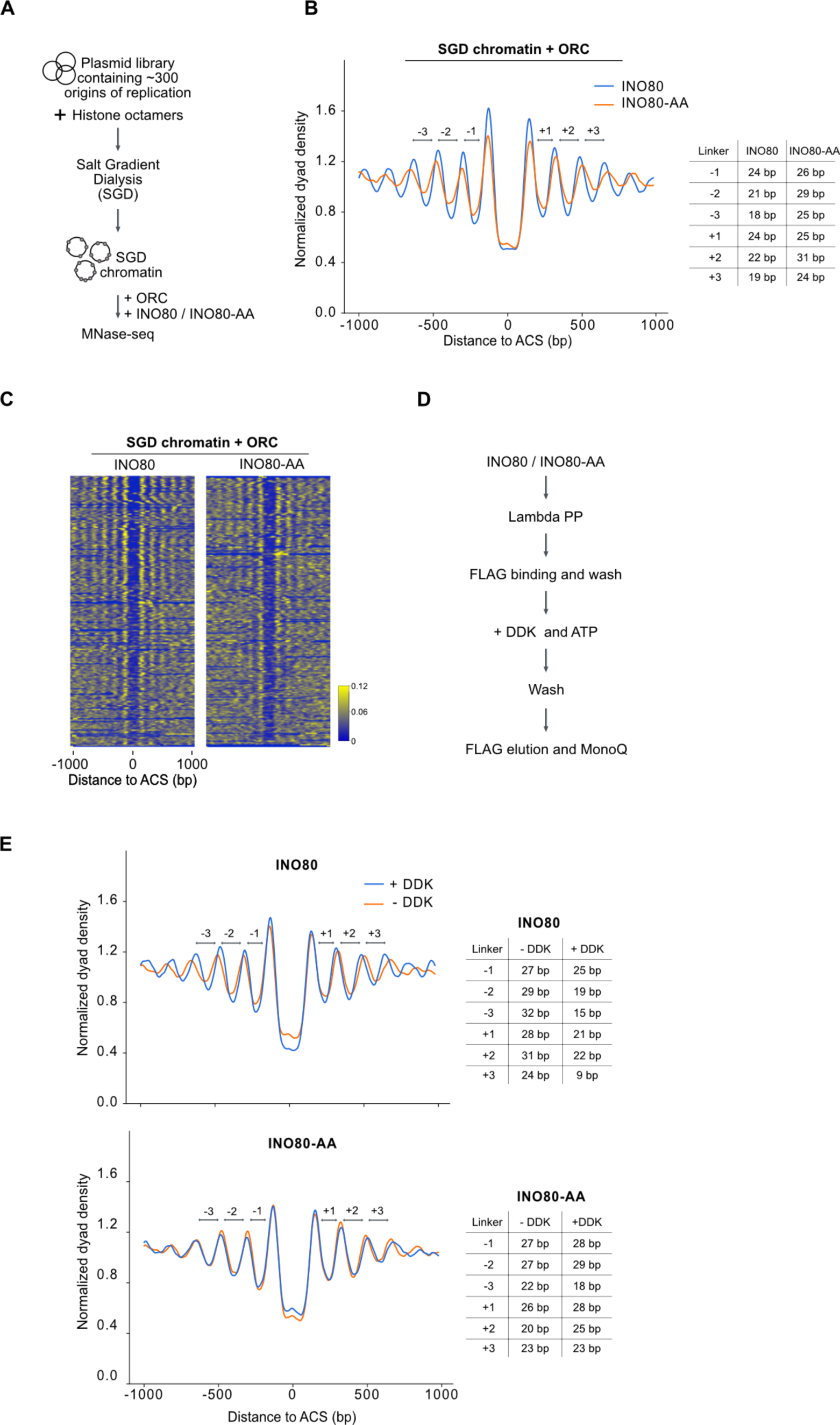
DDK-dependent phosphorylation is important for nucleosomal spacing activity of the INO80 complex. **A)** Outline of the genome-scale nucleosome reconstitution assay with purified INO80 and INO80-AA complexes to measure nucleosome spacing at origins of replication. **B)** Composite plot illustrating representative *in vitro* MNase-seq data for SGD chromatin incubated with ORC plus either INO80 (blue) or INO80-AA (orange) complexes as described previously (Chacin et al., 2023). Approximately 300 replication origins are aligned to the ORC binding motif ACS (ARS (Autonomously Replicating Sequence) Consensus Sequence). The accompanying table presents the linker lengths of the first, second, third linkers (DNA between nucleosomes) upstream and downstream of NFR (Nucleosome Free Region) generated by INO80 or INO80-AA complexes. N=2. **C)** Heat maps of the MNase seq data for the SGD chromatin presented in (B). Coverage was collected in 2-kb windows centered on ACS sites, smoothed using rolling averages with a window size of 25 bp and normalized to the square root. The heatmap was sorted from top to bottom according to pattern regularity. This regularity was defined by spectral density estimation over the entire 2-kb window at the frequency band corresponding to the nucleosomal repeat length in Saccharomyces cerevisiae (165 bp). **D)** Purified INO80 and INO80-AA complexes were dephosphorylated using Lambda Phosphatase (Lambda PP) and re-phosphorylated with purified DDK. N=2. **E)** Composite plot as in (B) illustrating representative *in vitro* MNase-seq data for SGD chromatin incubated with ORC plus either DDK phosphorylated (blue) or dephosphorylated (orange) INO80 and INO80-AA complexes. N=2.

To date, the results of our experiments indicate that DDK-dependent phosphorylation of the Arp8 subunit of INO80 plays a crucial role in maintaining its topological stability, which is closely linked to its spacing function. The next step was to analyse whether phosphorylation is important for INO80’s ability to space nucleosomes. Thus far, we have employed a mutant in which serine 65 and serine 233 were converted into alanines (residues that cannot be phosphorylated). This clearly demonstrates that both amino acids are important for INO80’s functionality. However, it does not provide evidence that phosphorylation is crucial for this. We decided against using a phospho-mimetic mutation (e.g. aspartic acid instead of serine or alanine), as we could clearly manipulate the phosphorylation status of INO80 by phosphorylation/dephosphorylation. Specifically, we dephosphorylated both INO80 wild-type and mutant complexes with lambda phosphatase (Lambda PP) in order to eliminate the majority of the phosphorylation events associated with the complexes. We then proceeded to treat or not treat the complexes with DDK, after which we re-purified the complexes. In the case of the INO80 wild-type complex, we observed a significant restoration of internucleosomal linker lengths after DDK phosphorylation (Figure 4E, left panel). In contrast, we did not observe this with the INO80-AA mutant complex, indicating that DDK phosphorylation is important for INO80’s ability to space nucleosomes.

### DDK-dependent phosphorylation of INO80 stimulates replication and is crucial during replication stress

Since all biochemical experiments have indicated that DDK plays a significant role in regulating INO80’s capacity to accurately space nucleosomes at replication origins, we proceeded to investigate the consequences of a loss of DDK-dependent phosphorylation of INO80 *in vivo*. Given INO80’s important roles in transcription, our initial aim was to determine whether DDK phosphorylation of INO80 might influence the transcriptome genome-wide. We postulated that a potential alteration in the expression of genes involved in the cell cycle (in particular those in G1 and S phase) or in replication genes might result in a defect in replication, thereby rendering conclusions about direct effects challenging. A comparison of the transcriptomes of isogenic *ARP8* wild-type and *arp8-AA* (in which serine 65 and serine 233 were converted into alanines) strains revealed no significant changes in the transcript levels of cell cycle or replication genes (Figure S2). The small proportion of misregulated genes was relatively limited and largely involved genes involved in metabolism (Figure S2). It is noteworthy that within this list, genes previously identified in a screen for transcriptionally misregulated genes under replication stress conditions were observed (Tkach et al., 2012) (Figure S2), which suggests that the *arp8* mutant may experience replication stress.

It has been demonstrated that replication problems can lead to recombinogenic DNA damage. In accordance with the hypothesis that DDK-dependent phosphorylation of INO80 plays a role in replication, we observed that the *arp8-AA* mutant exhibited a twofold increase in spontaneous levels of homologous recombination, as detected with a direct-repeat recombination assay (Figure 5A). We next assayed the ability of the *arp8-AA* mutant to cope with different forms of replication stress. The *arp8-AA* mutant exhibited a minor growth phenotype compared to *ARP8* wild type on full media without any drug supplementation (Figure 5B). A similar picture emerged when cells were grown on the methyl methanesulfonate (MMS), an agent inducing heat-sensitive DNA damage (Figure 5B). In stark contrast, *arp8-AA* mutants exhibited extreme sensitivity to HU (Figure 5B).

**Figure 5:**
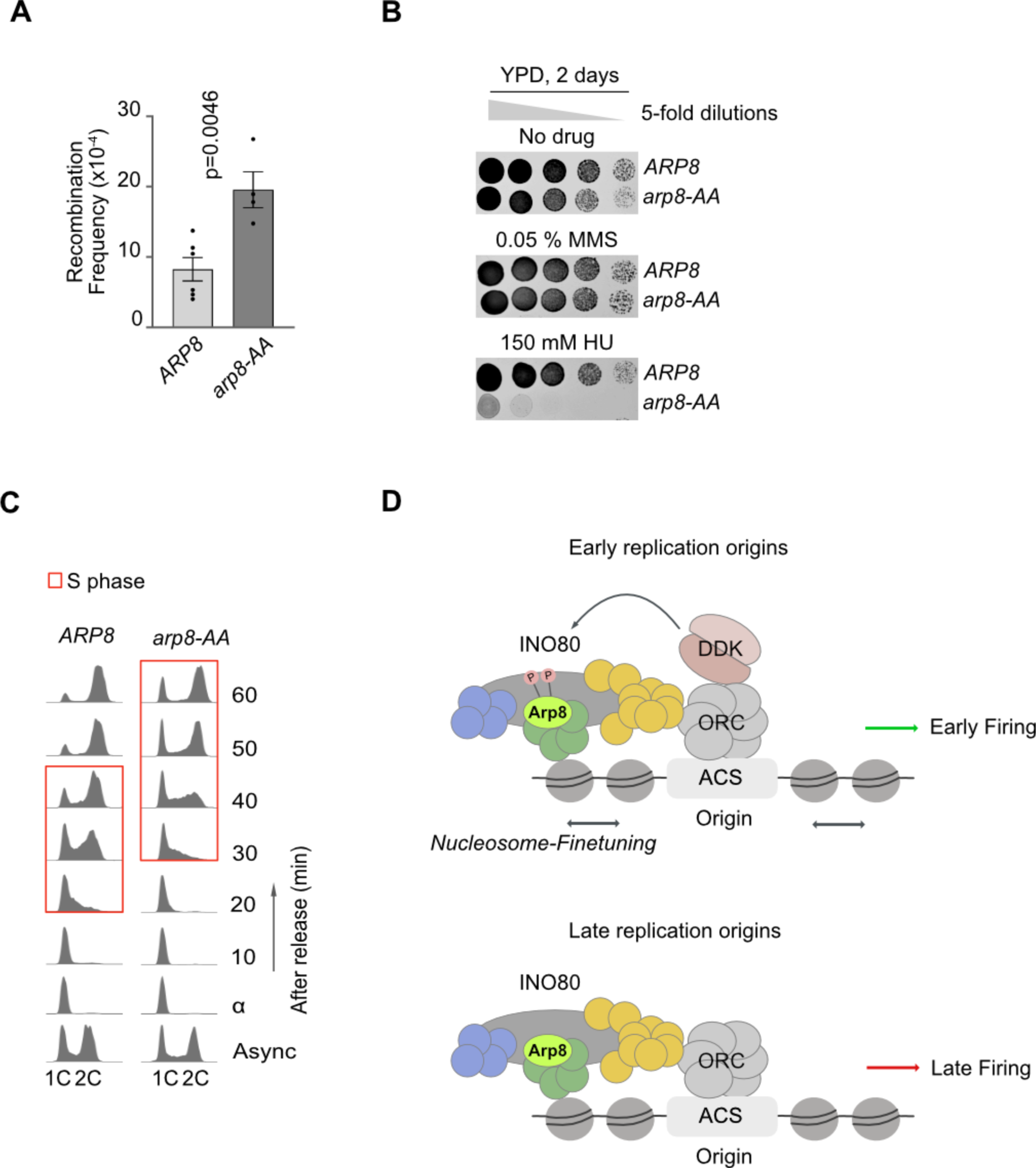
DDK-dependent phosphorylation of INO80 is important for efficient replication. **A)** Increase of recombination frequency in the *arp8-AA* mutant compared to *ARP8* WT. The graph is plotted with standard mean and error and p-values was obtained by using two tailed unpaired t-test calculation. N=4. **B)** Spot dilution growth assays (5-fold dilutions) with *ARP8* and *arp8-AA* strains. YPD: Yeast extract, Peptone, Dextrose (full medium); MMS: methyl methane sulphate and HU: hydroxyurea. N=2. **C)** Fluorescence Activated Cell-Sorting (FACS) profiles of *ARP8* versus *arp8-AA* cells before (async) and after synchronization in G1 phase with α-factor (α) and release (time points after release are indicated to the right). 1C and 2C indicate the position of non-replicated and replicated DNA fluorescence signal, respectively. Time points with cells in S phase are marked with a red rectangle. N=2. F) Model depicting the role of DDK activity at early replication origins, highlighting its function in fine-tuning INO80 activity through phosphorylation. See text for details.

The next objective was to determine whether the deficiencies in nucleosome positioning observed in our reconstitution experiments using the INO80-AA mutant complex were associated with defects in DNA synthesis *in vivo*. To this end, we monitored the progression of S phase in cells following pheromone-mediated cell cycle arrest or release, employing flow cytometry. The *arp8-AA* mutant exhibited a replication phenotype compared with the wild type, displaying a delay in the entry into S phase and a prolonged S phase (Figure 5C).

In conclusion, the combined results of our DDK screens yielded a network of approximately 400 phosphorylation events with high confidence. Within this network, we identified serine 65 and 233 of the Arp8 subunit of the INO80 chromatin remodeler as being phosphorylated by DDK. Further investigation, including structural, biochemical and cell biological studies, demonstrated that DDK-dependent phosphorylation of INO80 plays a role in regulating its activity to position nucleosomes at replication origins. Consequently, this is of significant importance for the replication process and the survival of cells under conditions of replication stress.

## Discussion

This study presents a global network of DDK-dependent phosphorylation events within the nucleus. A significant advancement of our approach is the utilisation of two independent methods to influence DDK activity (Figure 1A). Each method of inhibition has its own set of advantages and disadvantages. For instance, the elimination of DDK function through the use of a temperature-sensitive mutant represents a relatively clean approach that does not utilise chemicals such as kinase inhibitors, which can have a multitude of effects. However, the temperatures employed in the screen (25 °C for kinase “on” and 37 °C for kinase “shut off”) are not optimal for yeast. This increases the likelihood of false positives, as heat shock-associated phosphorylation events are more likely to occur at 37 °C. Conversely, there is a possibility that phosphorylation may not occur at 25 °C. Indeed, our observations indicated that serine 65 was particularly abundant when cells were grown at optimal temperature, thereby reinforcing the significance of our two-pronged screening pipeline. On the other hand, the activation of checkpoint kinases addresses the temperature issue, but checkpoint kinases such as Rad53, in addition influencing DDK binding to putative DDK targets, are likely to give rise to DDK-independent changes in phosphoproteomes. Consequently, we posit that our two-pronged approach is well-suited to the task of delineating a DDK-dependent phosphorylation network with a high degree of confidence.

Another advancement of our screening pipeline is the isolation of nuclei for the study of chromatin-associated processes. The presence of highly abundant cytosolic proteins may result in the masking of relatively low-abundant nuclear proteins, particularly chromatin factors, which may in turn result in a poor coverage of this class of proteins. Moreover, the phosphorylation sites on nuclear proteins may be challenging to associate with a specific kinase following whole-cell fractionation. For example, following cell disruption, a kinase that is typically excluded from the nucleus may gain access to and phosphorylate nuclear proteins in an unspecific manner, thereby significantly increasing the occurrence of false positives. The budding yeast is an exemplary system for this purpose, as it does not disintegrate its nuclear envelope during cell division, which enables the enrichment of nuclear fractions.

The focus of this study was the Arp8 subunit of INO80, which was identified within this network. Subsequent detailed follow-up studies identified a crucial role for DDK-dependent phosphorylation of INO80 in its ability to accurately position nucleosomes at replication origins. This was crucial for the initiation of replication and the survival of cells upon replication stress. Nucleosome architecture at replication origins is established during the G1 phase, prior to replication initiation (Chacin et al., 2023; Soriano et al., 2014). Our data are consistent with the finding that DDK interacts with ORC at replication origins in the G1 phase (Duncker et al., 2002; Pasero et al., 1999). It is noteworthy that DDK was demonstrated to be recruited to early-firing origins during the G1 phase by forkhead transcription factors Fkh1 and Fkh2 (Fang et al., 2017). This interaction was found to be crucial for origin function, possibly by recruiting replication factors from a competing and limiting pool. Our data now introduce a further layer of complexity, demonstrating that DDK directly targets a nucleosome spacing remodeler and that this influences replication efficiency (Figure 5D). We propose that this nucleosome “fine-tuning” mechanism may contribute to the RT programme and that this, *in vivo*, may preferably occur on early replicating origins. Pioneering studies have already identified a stimulatory role for spacing remodelers such as INO80 or ISW2 in replication *in vivo* (Papamichos-Chronakis et al., 2008; Vincent et al., 2008), the underlying mechanisms, however, remained obscure. More recent biochemical reconstitution studies have demonstrated that the actual spacing of nucleosomes is crucial for the process (Azmi et al., 2017; Chacin et al., 2023; Kurat et al., 2017). The observation that early origins exhibit better positioned nucleosomes compared to late origins is perfectly consistent with our model (Soriano et al., 2014). The precise influence of arrays of well-positioned nucleosomes on DNA-templated processes such as transcription or replication remains unknown. It has been postulated that closely packed nucleosomes in irregular arrays may impede transcriptional elongation (Ocampo et al., 2019). It seems plausible to suggest that a similar phenomenon may be occurring during replication. However, it is likely that there are additional factors at play, given that the lengths of lagging strands appear to correlate with nucleosome positioning (Smith et al., 2012). This suggests that regular arrays may play a crucial role in coordinating lagging and leading strand synthesis. It may therefore be the case that regular arrays are more important for replication than for transcription. In accordance with this hypothesis, the *arp8-AA* mutant, in which nucleosome positioning at origins was altered, resulting in replication defects (Figures 4 and 5), exhibited minimal changes in transcription (Figure S2).

In contrast to replication factors, which are solely involved in replication, chromatin factors are involved in a multitude of cellular processes, including transcription. The discovery that cell cycle kinases not only target core replication factors, but also chromatin factors involved in replication (Bansal et al., 2022; Chacin et al., 2021), may provide an explanation for how chromatin factors become “replication competent”. The precise mechanism by which phosphorylation of Arp8 renders INO80 replication competent remains unclear. It is conceivable that Arp8 phosphorylation is instrumental in the interaction with the barrier complex ORC at replication origins (Chacin et al., 2023). In the case of other barrier proteins, such as Abf1 or Reb1, which collaborate with INO80 to set up phased nucleosomal arrays at gene promoters (Krietenstein et al., 2016), it is possible that this process does not depend on Arp8 phosphorylation.

Collectively, our findings demonstrate that DDK plays a pivotal role in establishing the nucleosomal landscape at replication origins by targeting the nucleosome spacing remodeler INO80 (Figure 5D). Our findings provide further insight into the role of DDK at replication origins during the G1 phase. Our data suggest that accurate nucleosomal spacing is a key determinant in the decision of whether an origin fires early or late. The prevailing view of nucleosomes is no longer limited to their role as obstacles that must be overcome; rather, they are now regarded as gatekeepers of physiological DNA templated processes, such as replication.

---

**Figure S1:**
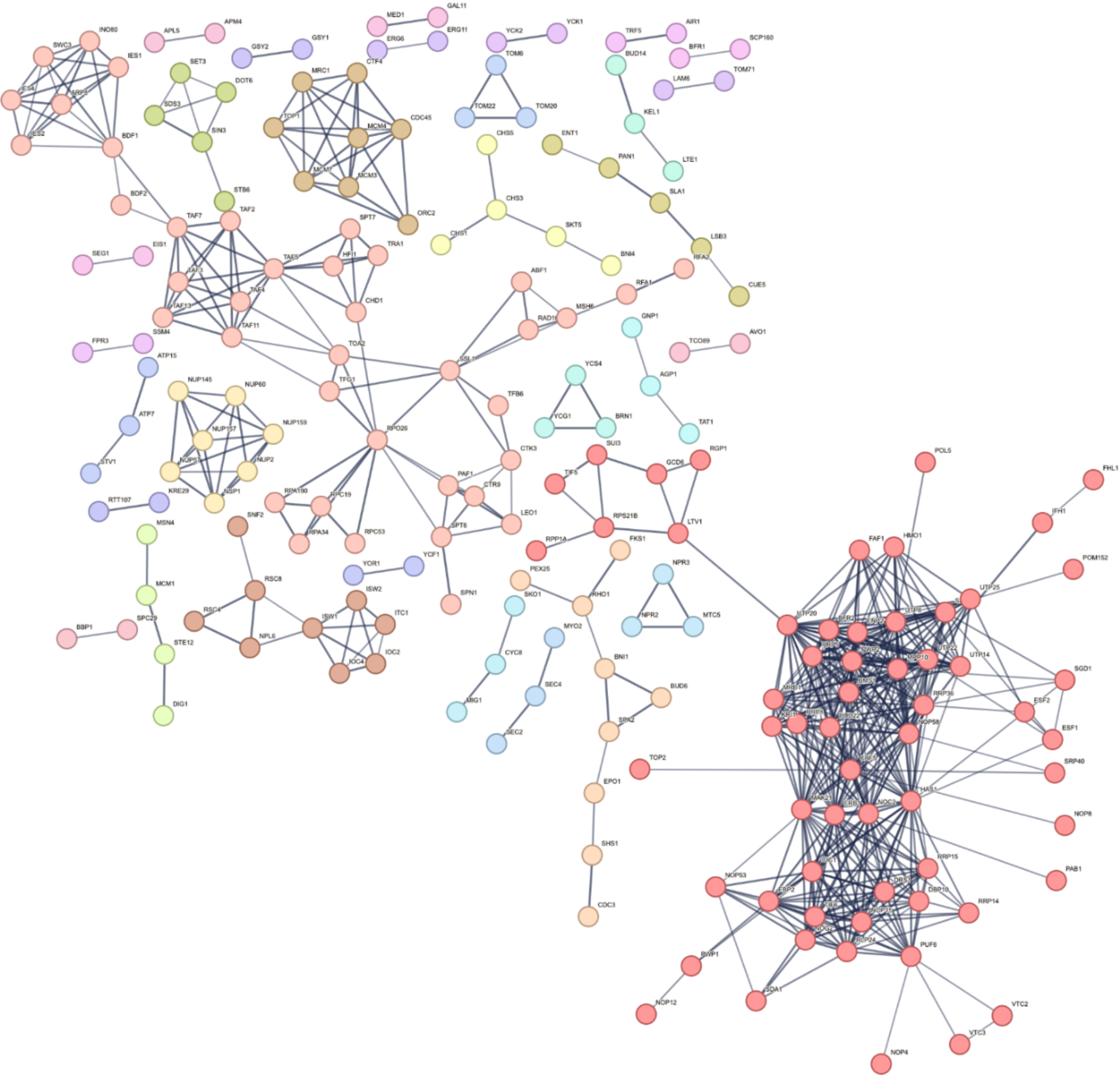
Network of putative DDK targets in complex involved in DNA replication and repair or chromatin remodeling and transcription. The network is part of a larger network constructed using data from the String Database (version 12.0. (Szklarczyk et al., 2023)) considering only high-quality interactions. The figure is generated using Cytoscape 3.10.2.

**Figure S2:**
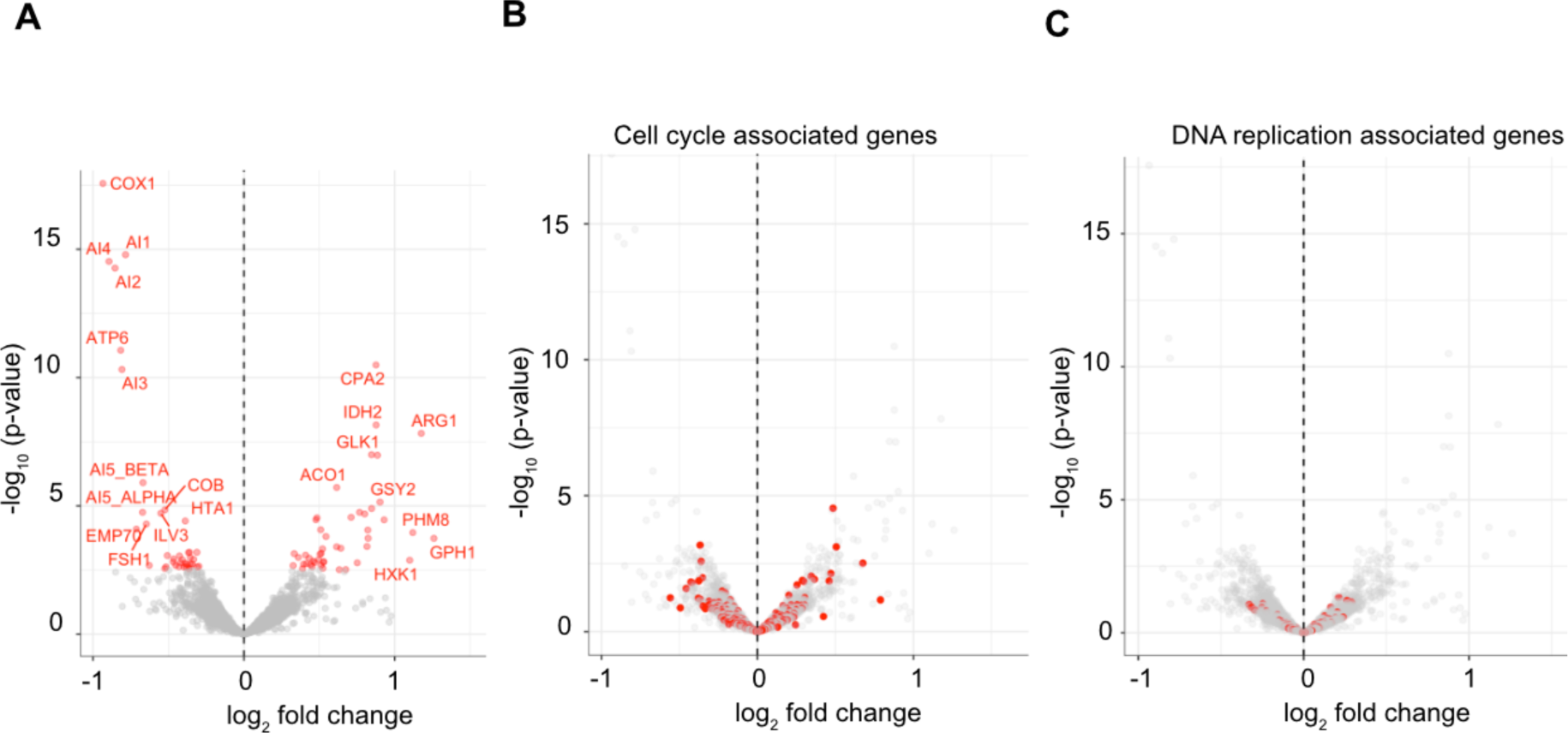
Transcriptional responses of the *arp8-AA* mutant **A)** Volcano plot representation of differential expression of genes in the *arp8-AA* versus the *ARP8*. Each dot on the plot represents an individual gene. Log2 fold change represents changes in gene expression that are upregulated or downregulated in *arp8-AA*. Red dots represent the genes, which are significantly up or down regulated. N=3. **B)** Volcano plot representation of differential expression of cell cycle associated genes in the *arp8-AA* versus the ARP8 WT. Red dots represents the genes, which are involved in the cell cycle process. **C)** Volcano plot representation of differential expression of DNA replication associated genes in the *arp8-AA* versus the ARP8 WT. Red dots represents the genes, which are involved in the DNA replication process.

## Acknowledgements

We would like to thank Mariia Likhodeeva, Karl-Peter Hopfner and Philipp Korber for discussing results. We also thank John Diffley for the *cdc7-4* stain and Helmut Blum and Stefan Krebs (LAFUGA) for high-throughput sequencing. This work was funded by the Deutsche Forschungsgemeinschaft (DFG) – the German Research Foundation – project ID 213249687 – SFB1064 to C.F.K and A. I. Work in the B.P. laboratory was supported by TU Dortmund University and DFG grants (466479039 and 445098914) awarded to B.P. The work in the A.A Laboratory was supported by the research funding to B.G.G by Programa Operativo FEDER 2014-2020 and Junta de Andalucía (project US-1380058). Work in the F.M.P. laboratory was supported by DFG (MU3613/1-2).

## Methods

### Strains and oligos

Yeast strains in this study were generated using standard genetic techniques. The strains were originated from the S288c genetic background. For the generation of INO80 expression strain, a 3X-FLAG was chromosomally inserted at the C-terminus of Ino80 in indicated strains using pBP83 as a template. All the oligonucleotides were synthetized from Merck. F and R stands for forward and reverse primers respectively.

A list of strains and primers used in this study can be found in Table 1 and Table 2.

**Table 1:**
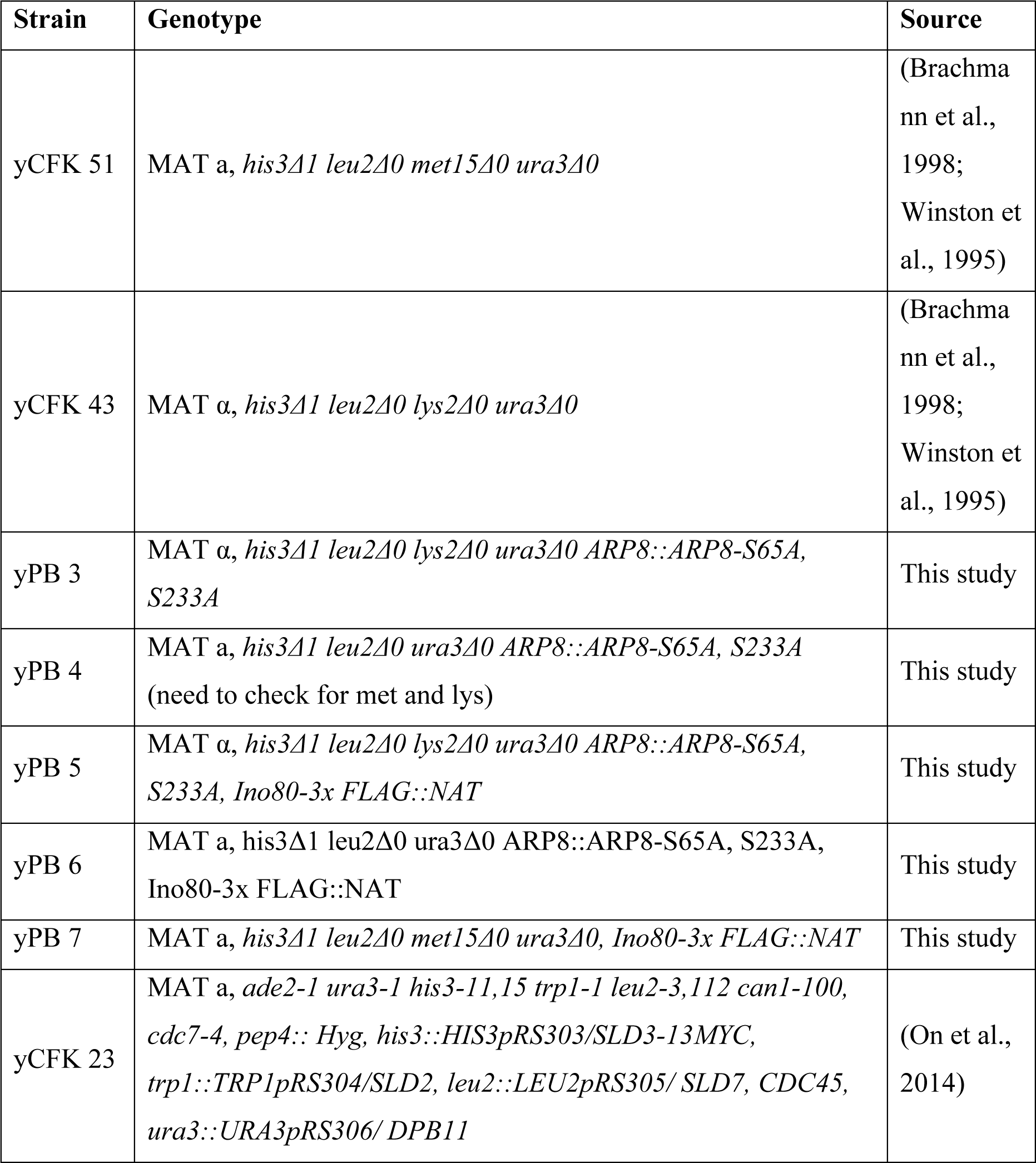
Yeast strains used in this study.

**Table 2:**
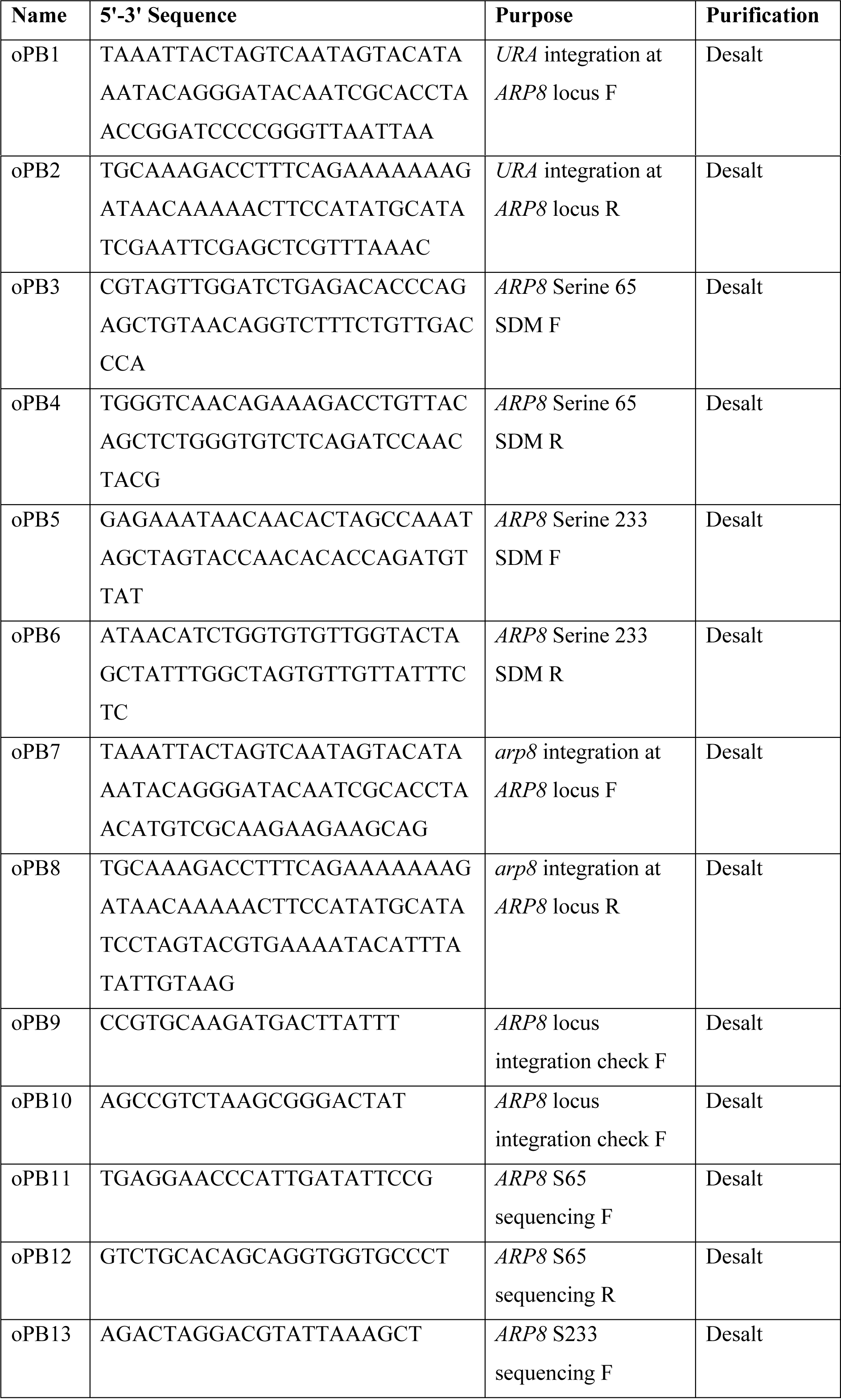
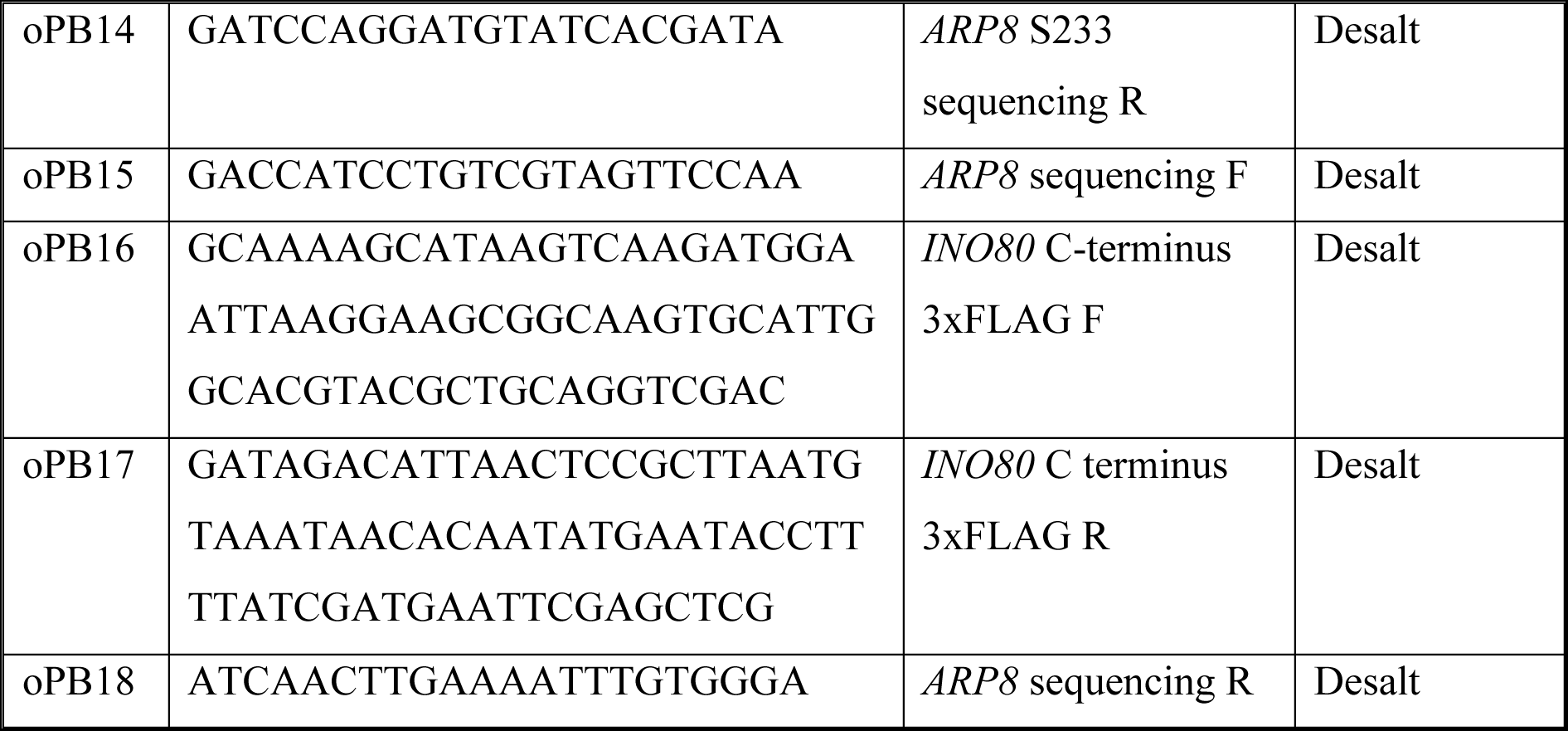
Primers used in this study.

### Protein expression and purification

The DDK and ORC complex were expressed and purified as previously described (Chacin et al., 2023; Kurat et al., 2017). INO80 expression and purification: The INO80 complex was expressed and purified as previously described before with modifications (Barrios et al., 2007; Biswas et al., 2005; Dyer et al., 2003; Kingston et al., 2011; Kurat et al., 2017; Shen, 2004). The INO80 complex was endogenously expressed and cells were grown in 16 litres YPD for 24 hours at 30° C. Cells were collected by centrifugation (6000 x g, 10 mins, 4° C) and pellet was resuspended in an equal volume of 2X lysis buffer with protease inhibitors 0.2 mM PMSF (Roth), 1 µM pepstatin A (Genaxxon), 1 µg/ml aprotinin (Genaxxon), and 2 µM leupeptin (Genaxxon) and . 1X lysis buffer is 25 mM HEPES-KOH pH 7.6, 500 mM KCl, 10% glycerol, 0.05% NP-40, 1 mM EDTA, 1 mM DTT, 4 mM MgCl2. Cells were frozen in liquid nitrogen in a dropwise manner and the frozen cells were crushed using a Freezer Mill (SPEX SamplePrep 6875 Freezer/Mill) (6 cycles for 2 min, crushing rate 15). To purify INO80, the frozen cell powder was thawed, resuspended in the 50 ml 1X lysis buffer with protease inhibitors. The insoluble material was cleared by ultracentrifugation ((235,000 x g, 1 h, 4° C). The supernatant was incubated with 1.5 ml pre-washed anti-FLAG M2 affinity gel resin (Sigma) in batch for 1h at 4 ° C. This was transferred into a disposable column, washed with 4 Column volume (CV) of lysis buffer and 1 CV of wash buffer (25 mM Tris-HCl pH 7.2, 200 mM KCl, 10% glycerol, 0.05% NP-40, 1 mM EDTA, 1 mM DTT, 4 mM MgCl2). INO80 was eluted by 1 ml of the same buffer with 0.5 mg/ml 3xFLAG peptide (Sigma), followed by 2X 1ml of buffer with 0.25 mg/ml 3xFLAG peptide. Eluted fractions were analysed by SDS-PAGE, pooled, and further purified with a Mono Q 5/50 GL column using a 15 CV gradient from 100 mM to 600 mM KCl (25 mM Tris-HCl pH 7.2, 5% glycerol, 0.05% NP-40, 1 mM EDTA, 1 mM DTT, 4 mM MgCl2). The fractions were analysed by SDS-PAGE, pooled, concentrated using a Vivaspin 20 (100 kDa MWCO Polyethersulfone, Merck). The Concentrated protein was then dialyzed for 1.5 hour in buffer containing 25 mM Tris-HCl pH 7.2, 600 mM NaCl, 40% glycerol, 1 mM EDTA and 1 mM DTT.

For cross-linking mass spectroscopy, the purified protein was equilibrated in 25 mM HEPES-NaOH pH 7.2, 300 mM NaCl, 5% glycerol, 1mM DTT, 4mM MgCl2 using Amicon Ultra Centrifugal Filter (3 kDa MWCO, Merck) for five times.

INO80-AA expression and purification: INO80-AA was purified using yPB5 which contains two point mutations and 3X-FLAG at the C-terminus of Ino80 subunit. For the protein expression, the INO80 complex was endogenously expressed and cells were grown in 24 litres YPD for 24 hours at 30° C, collected, processed and purified similar to INO80.

To dephosphorylate INO80 and INO80-AA, purified complexes were treated with Lambda phosphatase (NEB) at a concentration of 5 µL per ml protein sample for a period of 2 hours at 16 °C. This was followed by the addition of alkaline phosphatase at a concentration of 5 µL per 1 ml of protein sample for a period of 6 hours at 4 °C. The phosphatases were then removed by rebinding the complexes to FLAG resin and washing them as described above. The complexes were either treated or not treated with DDK and ATP on the beads for one hour at 16 °C. DDK was removed by washing as described above. Subsequently, the complexes were eluted and the FLAG peptide was removed by MonoQ purification.

### Cell cycle arrest and release

Cells were grown at 30° C in YPD to OD 600nm = 0.2 - 0.5 (OD was measured in Thermo Scientific GENESYS 20 spectrophotometer) and then arrested in G1 with alpha-factor (GenScript (RP01002)– 10 mg/ml stock concentration and 10 µg/ml working concentration) for 120 mins in total. A second dose of alpha-factor was added after 60 mins. Cells were then washed with pre-warmed YPD twice and released into fresh pre-warmed YPD to resume the cell cycle. Samples for FACS analyses and mass spectrometry were taken at indicated time points and processed according to standard protocols.

### Flow cytometry

Cell cycle arrest and release samples were collected as described above. For flow cytometry, samples were processed as described previously (Reusswig et al., 2022). Briefly, yeast cells were harvested by centrifugation, resuspended in 50 mM Tris-HCl pH 8.0 / 70% ethanol and stored at 4 ° C for minimum one hour for permeabilization and fixation. Cells were then washed once with 50 mM Tris-HCl, pH 8.0 and treated with 200 *μ*g RNase A (Sigma, R4875; diluted in 10 mM Tris-HCl, pH 7.5, 10 mM MgCl2) for at least 4 h at 37 °C, and subsequently treated with 400 *μ*g Proteinase K (Sigma, P2308; diluted in 10 mM Tris-HCl, pH 7.5) for 30 min at 50 °C. Cells were later resuspended in 50 mM Tris-HCl, pH 8.0, sonicated and diluted 1:20 with 50 mM Tris-HCl, pH 8.0 containing 0.5 *μ*M SYTOX Green (Invitrogen, S7020) and measured with a MACSquant Analyzer Flow Cytometer (Milteny Biotec).

### *In vitro* DDK kinase reactions

15 nM of WT INO80 and mutants, were incubated with 5 nM DDK in buffer containing 100 mM potassium-glutamate, 25 mM HEPES-KOH pH 7.6, 10 mM Mg(OAc)2, 0.02% NP-40, 1 mM DTT, 10 mM ATP and 5 mCi ^32^P-g-ATP for 30 mins at 30 ° C. Proteins were then separated on SDS-PAGE, gel were dried, exposed with Super RX Medical X-Ray Film (FUJI) and autoradiograph was developed using a Typhoon scanner phospho imager (GE Healthcare).

### Genetic analysis of recombination

Recombination frequencies were calculated by transforming the yeast strains with the pRS316-L plasmid containing two truncated repeats of the *LEU2* gene sharing 600 bp of homology and placed on a mono-copy *CEN-*based plasmid (Gómez-González et al., 2011). Recombinants were obtained by plating appropriate dilutions in SC media lacking leucine and uracil. To calculate total number of cells, they were plated in the same media supplemented with leucine. All plates were grown for 3-7 days at 30 ° C. For each transformant, the median value of six independent colonies was obtained. The mean and SEM of at least three independent experiments performed with independent transformants was plotted.

### Spot dilution assay

Cells were grown overnight to near saturation and OD 600 nm was measured in technical replicates after 1:10 dilutions in water. Cells were diluted to OD 600nm = 1.0 in 250 µl water and 5-fold dilutions were generated. Seven microliters of the dilutions were spotted on YPD plates and YPD plates supplemented with desired drug as necessary. Plates were incubated at 30 ° C for 2 days.

### Transcriptome analysis

Cells were generally grown overnight to OD 600nm = 0.8 - 1.0 in 10 ml YPD complete media. Next day, cells were reinoculated in 100ml fresh YPAD media at 0.1 OD. At OD 600 nm = 0.4 cells were collected by centrifugation at 4500 x g, 10 mins, 4° C, the pellets were washed once with cold distilled water, flash frozen in liquid nitrogen and stored at -80 °C.

For extraction of total RNA, pellets were resuspended in 1 ml QIAzol Lysis Reagent (Qiagen) and mixed with 250 µl of zirconia beads (Roth). Cells were lysed using the Precellys 24 homogenizer (Bertin) for 3 x 30 sec with 5 min rest in between on ice, followed by centrifugation at 12000 x g, 10 min, 4° C. To the cleared lysate, 200 µl chloroform was added and mixed by brief vortexing for 15 sec. Samples were incubated at room temperature (RT) for 10 min followed by centrifugation at 12000 x g, 10 min at 4° C. The aqueous phase was extracted by adding 500 µl chloroform followed by brief vortexing and centrifuged at 12000 x g for 10 min at 4° C. To the aqueous phase, 500 µl isopropanol was added for 15 min at 4° C and pelleted by centrifugation at 12000 x g, 10 min, 4° C to precipitate RNA. The RNA pellets were washed two times with 1 ml of 75% ethanol. The RNA pellet was dissolved in 150 µl RNase-free water for 30 min at 55° C and quantified photometrically with DeNovix Spectrophotometer. For each sample, 10 µg RNA was treated with 10 units recombinant DNase I (Roche) for 1 h at 37 ° C, purified with Agencourt RNAClean XP beads (Beckman Coulter) and quantified photometrically with DeNovix Spectrophotometer.

Library preparation and sequencing: For each sample, 1 µg DNaseI-treated RNA was used for rRNA depletion and library preparation. Ribosomal RNA was depleted using the *S. cerevisiae* - specific riboPOOL kit (siTOOLs Biotech) according to the manufacturer’s protocol. rRNA-depleted RNA was purified with Agencourt RNAClean XP beads (Beckman Coulter) and analyzed on 4150 TapeStation System using a High Sensitivity RNA Screen Tape (Agilent). Of the rRNA-depleted samples directional libraries were prepared using the NEBNext Ultra II Directional RNA Library Prep Kit for Illumina (New England Biolabs) in accordance to the recommended protocol. The quality of the libraries was assessed on 4150 TapeStation System using a High Sensitivity D1000 Screen Tape. Libraries were sequenced on an Illumina NextSeq 2000 instrument in paired-end mode.

Data analysis: Sequencing reads were pseudoaligned to the yeast transcriptome (Ensembl build R64, annotation version 108) using kallisto (version 0.48) using default parameters. Data was further processed in R/bioconductor. Differential expression was tested with DESeq2(version 1.36.0) using the experimental batch as random factor.

### Nuclei isolation

Yeast nuclei were isolated as described previously with modifications (Almer et al., 1986). Yeast cells were grown overnight to OD 600 nm = 0.4 – 0.6 in 200 or 500 ml YPD complete media. Cells were harvested by centrifugation at 4500 x g, 10 min, 4° C and the pellets were washed once with cold distilled water. The weight for washed pellet (wet weight) was determined and resuspended in 2x wet weight of preincubation solution containing 0.7 M β - mercaptoethanol and 2.8 mM EDTA pH 8.0. Cells were shaken for 25 - 30 mins at 30° C, then washed with 40 ml of cold 1 M sorbitol, and finally the pellet was resuspended 5 ml buffer (5 mM β -mercaptoethanol in 1 M sorbitol) per gm wet weight of the pellet. OD 600 nm of resuspended pellet was determined using 1:100 dilution in water. To digest the cell wall (spheroblasting), Zymolyase was freshly dissolved in water and was then added to the resuspended pellet. 2 mg Zymolyase was added per gram of wet weight and incubated at 3° C for 20 - 30 min. The cell wall was digested until the absorbance at 600 nm was decreased to 80 - 90% of the starting OD when the digestion was considered to be complete. Spheroblasts were harvested by centrifugation (2500 x g, 5 min, 4° C), washed one time with 40 ml cold 1 M sorbitol and resuspended in a Ficoll buffer (18% Ficoll type 400 (Sigma), 20 mM KH2PO4 pH 6.8, 1 mM MgCl2, 0.25 mM EGTA pH 8.0, 0.25 mM EDTA pH 8.0). 7 ml of the Ficoll buffer was added per gram weight of cells. Lastly, nuclei were aliquoted to desired wet weight (0.5 or 1 g) and centrifuged at 12000 x g, 30 mins, 4° C. The nuclei pellets were frozen in dry ice/ethanol bath and were stored at -80 ° C until further processing.

### Nuclei processing for mass spectrometry

Mass spectrometry samples for label free phospho proteome and whole proteome analysis were processed exactly as previously described (Humphrey et al., 2018). All reagents were prepared with mass spectrometry grade reagents. Briefly, thawed nuclei were first washed with 100 mM Tris-HCl pH 8.5 to remove phosphate buffer residues. Chilled lysis buffer containing 4% Sodium deoxycholate (Sigma) and 100 mM Tris-HCl pH 8.5 was added such that total volume is about 600 μl. The lysates were heat inactivated at 95° C for 5 mins and then homogenized by bath-sonication using Biorupter Pico at 4° C (two times 10 cycles each of 30 sec on and 30 sec off at maximum output power). The cleared lysates were collected in a new eppendorf tube by centrifugation and protein concentration was determined using Bicinchoninic acid (BCA) protein assay kit (Pierce). All samples were diluted to a final concentration of 300 μg in 270 μl of lysis buffer. Reduction/alkylation buffer containing 100 mM Tris(2-carboxyethyl) phosphine hydrochloride (TCEP) (Sigma) and 400 mM of 2-chloroacetamide (CAM) (Sigma) was added at 1:10 ratio of the total volume (30 μl) and incubated for 5 mins, 45 °C with shaking at 1500 rpm. At room temperature, using 1:100 enzyme to substrate ratio, 3 μg each of Lys-C and trypsin was added to each 300 μg sample and digested overnight at 37 ° C with shaking at 1500 rpm.

Next day the digested peptides were separated such that 2/3 of the sample was processed for phospho proteome and 1/3 was processed for whole proteome. Stage tips were prepared as described earlier (Rappsilber et al., 2007).

For phospho proteome, phospho peptides were enriched by TiO2 based enrichment and eluted using C8 material (Empore) on C8 stage tips. Collected phopsho peptides were desalted and eluted using SDB-RPS (Styroldivinylbenzol Reversed Phase Sulfonat) material (Empore) on SDB-RPS stage tip. The eluate was evaporated under vacuum to dryness at 45° C. Phospho peptides were then reconstituted in 10 μl of MS loading buffer containing 0.3% Trifluoroactetic acid (TFA)(Merck) and 2% Acetonitrile (ACN) (Roth).

For whole proteome, 1/3 of the digested peptides were dissolved with 200 μl isopropanol and 1% TFA. Peptides were desalted and eluted using by SDB-RPS material on SDB-RPS stage tip. The eluate was evaporated under vacuum to dryness at 45° C and peptides were reconstituted in 15 μl of MS loading buffer.

### Mass spectrometric measurements and data analysis

For LC-MS purposes, desalted peptides were injected in an Ultimate 3000 RSLCnano system (Thermo Fisher Scientific) and separated in a 25-cm analytical column (75µm ID, 1.6µm C18, IonOpticks) with a 50 min gradient from 2 to 35% or 60-min gradient from 2 to 32% ACN in 0.1% formic acid for proteome and phosphoproteome analysis, respectively.

Phospho proteome: The effluent from the HPLC was directly electrosprayed into an Orbitrap Exploris-480 (Thermo Fisher Scientific) operated in data dependent mode to automatically switch between full scan MS and MS/MS acquisition. Survey full scan MS spectra (from m/z 350–1400) were acquired with resolution R=60,000 at 400 m/z (AGC target of 3×10^6^). The 15 most intense peptide ions with charge states between 2 and 5 were sequentially isolated to a target value of 2×10^5^, fragmented at 30% normalized collision energy and acquired with resolution R=15,000. Typical mass spectrometric conditions were: spray voltage, 1.5 kV; no sheath and auxiliary gas flow; heated capillary temperature, 275° C; ion selection threshold, 5×10^3^ counts. For phosphopeptides, MS2 resolution was increased to R=30,000 and ion selection threshold to 3×10^4^ counts.

Whole proteome: Eluting peptides were ionized in a nanoESI source and on-line detected on a QExactive HF mass spectrometer (Thermo Fisher Scientific). The mass spectrometer was operated in a TOP10 method in positive ionization mode, detecting eluting peptide ions in the m/z range from 375 to 1,600 and performing MS/MS analysis of up to 10 precursor ions. Peptide ion masses were acquired at a resolution of 60,000 (at 200 m/z). High-energy collision induced dissociation (HCD) MS/MS spectra were acquired at a resolution of 15,000 (at 200 m/z). All mass spectra were internally calibrated to lock masses from ambient siloxanes. Precursors were selected based on their intensity from all signals with a charge state from 2+ to 5+, isolated in a 2 m/z window and fragmented using a normalized collision energy of 27%. To prevent repeated fragmentation of the same peptide ion, dynamic exclusion was set to 20 s.

MaxQuant search parameters: For phospho proteome identification, MaxQuant 2.0.3.0 software package was used. Parent ion and fragment mass tolerances were 8 ppm and 0.5 Da, respectively, and allowance for two missed cleavages was made. Yeast canonical protein database from UniProt (*Saccharomyces cerevisiae* (strain ATCC 204508 / S288c)), filtered to retain only the reviewed entries were used for the searches. Regular MaxQuant conditions were the following: site FDR, 0.01; protein FDR, 0.05; minimum peptide length, 6; variable modifications, oxidation (M); phospho (STY); fixed modifications, carbamidomethyl (C); peptides for protein quantitation, razor and unique; minimum peptides, 2; minimum ratio count, 2. Proteins were validated on the basis of at least one unique peptide detected in the proteome of all the three replicates or in at least two of the three replicates.

For whole proteome identification, MaxQuant search parameters were identical except for the variable modifications, which were oxidation (M); acetyl (protein N-term); acetyl (K); dimethyl (KR); methyl (KR).

Data analysis: The phospho-proteomics data was analysed using an R-script developed in-house. Differential and quantitative analysis was performed using phospho-sites with a 75% or higher probability of occurrence (according to MaxQuant output). Phospho-sites and proteins that were present in at least 75% of the replicates were considered for the downstream analysis. Intensity based absolute quantification (iBAQ) values were used to quantify the abundance of phospho-sites and compare it in different conditions. Differential expression analysis at the whole and phospho proteome level was carried out using the DEP package. Briefly, after filtering for all the experimental and analytical contaminants missing values were imputed by the Bayesian principal component analysis (BPCA) method followed by limma statistical analysis (Zhang et al., 2018) using a p-value cut-off of 0.05. The phospho-site abundances were normalised to the total protein abundance of the respective proteins.

GO term analysis: GO term analysis for the corresponding proteins of the overlapping phosphosites within the DDK active fractions of sc7-4 and Hydroxyurea screens were performed in ImShot (Aftab et al., 2022) Pusing the following parameters: p-value cutoff – 0.05, p-value adjustment – Benjamini-Hochberg (BH), Database: org.Sc.sgd.db, Redundancy removed, Minimal and maximal size of genes – 1 and 500 respectively.

### Cross-linking mass spectrometry

Firstly, the equilibrated purified protein was titrated against different DSBU (Thermo Fischer Scientific) cross-linker concentration to determine the optimal cross-linking concentration for each protein. The INO80 complex (1 µg in total of INO80 or INO80-AA) was cross-linked for 20 min at 30° C on a thermomixer at 1200 rpm. Cross-linked product was separated on SDS-PAGE followed by silver staining. The optimal INO80 concentration was determined be 25 µg and 15 µg at 1200 rpm, 20 min, 30° C. The optimal DSBU cross-linker concentration for INO80 was 58 µM and for the INO80-AA was 100 µM.

The reaction with optimal protein and cross-liner concentration was performed at 1200 rpm, 20 min, 30° C. The reaction was quenched by adding ammonium bicarbonate to a final concentration of 100 mM and incubated for 10 min at 30°C subsequently followed by protein denaturation, alkylation, and tryptic digest. Cross-linked samples were denatured by adding two sample volumes of 8 M urea, reduced with 5 mM Tris (2-carboxyethyl) phosphine (TCEP) and alkylated by the addition of 10 mM iodoacetamide (IAM) for 40 min at RT in the dark. Proteins were digested with 0.5 μg Lys-C at 35°C for 2 hr, diluted with 50 mM ammonium bicarbonate, and digested with trypsin 0.5 μg overnight. Peptides were acidified with 1% trifluoroacetic acid (TFA) and purified by reversed phase chromatography using C18 material (Empore) on C18 stage tips. Further, cross-linked peptides were enriched on a Superdex Peptide PC 3.2/30 column using water/acetonitrile/TFA (75/25/0.1) as mobile phase at a flow rate of 50 μl/min. Fractions containing cross-linked peptides were analysed by liquid chromatography (Dionex 3000, Thermo Fisher Scientific) coupled to tandem mass spectrometry (LC-MS/MS) using a TimsTOF Pro instrument (Bruker Daltonics).

### LC-MS Analysis of enriched cross-linked peptides

Cross-linked peptides were injected and separated on a PepSep column (25 cm, inner diameter 150 µm, Bruker Daltonics) by an online reversed-phase chromatography with a 50 min gradient from 3 to 43 % of Buffer B containing 100% ACN, 0.1% formic acid) at a flow rate of 300 nl/min. Eluting peptides were directly sprayed through the CSI source into the TimsTOF Pro. Each sample was measured in three independent technical replicates. The mass spectrometric measurement was performed in data-dependent acquisition mode with a top 10 method. The same settings were applied as described (Ihling et al., 2021). As template the standard DDA-PASEF MS Method provided by Bruker Daltonics was used. Only precursor ions of +3 to +8 charge (in case for DSBU +2 to +8 charge) were selected for fragmentation scan. Raw data files were searched against MaxQuant software package (version 2.0.2.0). Following changes have been applied for the MaxQuant search: Enzyme specificity set to trypsin with a maximum number of missed cleavages 3; DSBU specificity linking (K, S, T, Y); fixed modifications carbamidomethyl (C); variable modifications, oxidation (M). PSM FDR crosslink set to 5%. Inter- and Intra-Crosslinks were filtered by applying an MS1 tolerance window of -3 to 3 ppm and a score ≥ 60 (Yılmaz et al., 2022). Cross-links were visualized as network plots using the webserver xiNET (Combe et al., 2015).

### Salt gradient dialysis (SGD) chromatin

Chromatin in this study was assembled using salt gradient dialysis (SGD) as previously described with modifications (Chacin et al., 2023; Ludwigsen et al., 2018).

Chromatin used for ATP hydrolysis assays was prepared as follows. A DNA fragment containing 25 consecutive copies of a modified 197-bp Widom-601 nucleosome positioning sequence was cut out of the plasmid pFMP232 using appropriate restriction enzymes (Ludwigsen et al., 2018). All steps were performed at 4 ° C. Briefly, histone octamers and 0.1 µg/µl digested plasmid DNA are mixed in 100 µl buffer containing 10 mM Tris–HCl pH 7.6, 2 M NaCl, 1 mM EDTA pH 8, 0.01% NP-40, 1 mM DTT. Samples were placed in Slide-ALyzer devices (MWCO 7K, Thermo Fisher Scientific) in 1 L high salt buffer containing 10 mM Tris–HCl pH 7.6, 2 M NaCl, 1 mM EDTA pH 8, 0.01% NP-40, 1 mM DTT. This was dialyzed in 3 L of low salt buffer containing 10 mM Tris–HCl pH 7.6, 50 mM NaCl, 1 mM EDTA pH 8, 0.01% NP-40, 1 mM DTT into the beaker over a time period of at least 24h at flow rate of ∼ 2 ml / min using peristaltic pump and tubing. Chromatin was then dialyzed for 2 h with 1 L low salt buffer at 30 °C and stored at 4 ° C.

For Nucleosome positioning assay, a yeast origin plasmid library containing ∼300 ARS (OriDB) sequence was generated from the *S. cerevisiae* genomic library (Open Biosytems).

Selected plasmids contained an origin of replication at least 1000bp away from the border of plasmid backbone and yeast genomic insert. Using SGD, 10 µg of origin plasmid library DNA was combined with Drosophila embryo histone in 100 µl SGD buffer containing 10 mM Tris– HCl pH 7.6, 2 M NaCl, 1 mM EDTA pH 8, 20 µg BSA, 0.05% Igepal to a saturated assembly degree. Samples were placed in Slide-ALyzer devices (Thermo Fisher Scientifc) in 300 ml high salt buffer containing 10 mM Tris–HCl pH 7.6, 2 M NaCl, 1 mM EDTA pH 8, 0.05% Igepal, 14.3 mM β-mercaptoethanol). This was dialyzed in 3 L of low salt buffer containing 10 mM Tris–HCl pH 7.6, 50 mM NaCl, 1 mM EDTA pH 8, 0.05% NP-40, 1.4 mM β-mercaptoethanol using peristaltic pump at 7.5 rpm, 30 ° C for 16 hours. Chromatin was then dialyzed for 1 h with 1 L low salt buffer at 30 ° C and stored at 4 ° C.

### ATP hydrolysis coupled with NADH oxidation assay

30 nM of the purified INO80 and INO80-AA, were incubated in buffer containing 100 mM potassium-glutamate, 25 mM HEPES-KOH pH 7.6, 10 mM Mg(OAc)2, and 1 mM DTT supplemented with 3 mM ATP (Sigma) and 3 mM MgCl2, 0.6 mM NADH (Sigma), 3 mM phosphoenolpyruvate (PEP) (Molecula) and 16 U/ml lactic dehydrogenase/pyruvate kinase enzymes (Sigma). 24 µl reactions were prepared in 384 well plates (Greiner) in buffer without ATP and MgCl2. The reactions were tested in sets of 3 technical replicates with either 100 ng/µl DNA (pFMP232) or 90 nM, 25mer nucleosome arrays assembled on the 25 Widom-601 repeats in pFMP232. To confirm saturation of ATP binding, reactions were also tested in sets of three technical replicates with either 300 ng/µl DNA (pFMP232) or 270 nM, 25mer chromatin originating from pFMP232.

ATPase reactions were initiated by adding an ATP / MgCl2 mix to a final concentration of 3 mM each. Plates were incubated at 26 ° C in a Biotek PowerWave HT plate reader for 60 mins and absorption at 340 nm was determined every 15 s. For analysis, each time course was fitted to a linear function within a time range where all reactions were linear. From the slope of the reaction and the extinction coefficient of NADH (6220 M^−1^ cm^−1^), the change in NADH concentration was calculated (for 30 μL reactions in Greiner plates, the path length was 0.27273 cm). As oxidation of one NADH equals the hydrolysis of one1 ATP, the ATP hydrolysis rates were calculated from the slopes.

### Nucleosome positioning assay

The assay was performed with 30nM ORC and 20nM INO80 or INO80-AA in a buffer containing 20 mM HEPES pH 7.5, 50 mM NaCl, 3 mM MgCl2, 2.5 mM ATP, 2.5 mM DTT, 0.5 mM EGTA pH 8, 12% glycerol, and started by the addition of SGD chromatin, incubated for 2 hours at 30 ° C and stopped by addition of 0.2 U apyrase (NEB), incubated at 30 ° C for 20 mins. In order to generate mostly mononucleosomal DNA, the reaction was incubated with 100 U MNase (Sigma-Aldrich) and 1.5 mM CaCl2 for 5 min at 30 ° C. The digestion was stopped by the addition of 10 mM EDTA and 0.5% SDS followed by a proteinase K treatment for 1 hour at 37 ° C and ethanol precipitation. Samples were run in 1.5% agarose gels for 1.5h at 110 V constant in 1X TAE (40 mM Tris, 20 mM acetic acid, 1 mM EDTA), and mononucleosomal DNA was excised and purified using DNA purification kit.

The sequencing libraries were prepared using 10–50 ng mononucleosomal DNA. The samples were diluted to 10 nM, pooled according to the sequencing reads required (∼5 million reads per sample), and quantified via BioAnalyzer (Agilent). The pool was sequenced with an Illumina NextSeq 1000 in 60 bp paired-end mode (Laboratory for Functional Genome Analysis, Ludwig-Maximilians-Universität Munich). The MNase-seq data was analysed as previously described (Chacin et al., 2023).

## References

Abd Wahab, S., & Remus, D. (2020). Antagonistic control of DDK binding to licensed replication origins by Mcm2 and Rad53. Elife, 9. doi:10.7554/eLife.58571

Akerman, I., Kasaai, B., Bazarova, A., Sang, P. B., Peiffer, I., Artufel, M., … Mechali, M. (2020). A predictable conserved DNA base composition signature defines human core DNA replication origins. Nat Commun, 11(1), 4826. doi:10.1038/s41467-020-18527-0

Argunhan, B., Leung, W. K., Afshar, N., Terentyev, Y., Subramanian, V. V., Murayama, Y., … Tsubouchi, H. (2017). Fundamental cell cycle kinases collaborate to ensure timely destruction of the synaptonemal complex during meiosis. Embo j, 36(17), 2488–2509. doi:10.15252/embj.201695895

Azmi, I. F., Watanabe, S., Maloney, M. F., Kang, S., Belsky, J. A., MacAlpine, D. M., … Bell, S. P. (2017). Nucleosomes influence multiple steps during replication initiation. Elife, 6. doi:10.7554/eLife.22512

Bansal, P., & Kurat, C. F. (2022). Yta7, a chromatin segregase regulated by the cell cycle machinery. Mol Cell Oncol, 9(1), 2039577. doi:10.1080/23723556.2022.2039577

Bell, S. P., & Labib, K. (2016). Chromosome Duplication in Saccharomyces cerevisiae. Genetics, 203(3), 1027–1067. doi:10.1534/genetics.115.186452

Berbenetz, N. M., Nislow, C., & Brown, G. W. (2010). Diversity of eukaryotic DNA replication origins revealed by genome-wide analysis of chromatin structure. PLoS Genet, 6(9), e1001092. doi:10.1371/journal.pgen.1001092

Bloom, J., & Cross, F. R. (2007). Multiple levels of cyclin specificity in cell-cycle control. Nat Rev Mol Cell Biol, 8(2), 149–160. doi:10.1038/nrm2105

Bousset, K., & Diffley, J. F. (1998). The Cdc7 protein kinase is required for origin firing during S phase. Genes Dev, 12(4), 480–490. doi:10.1101/gad.12.4.480

Brahma, S., Ngubo, M., Paul, S., Udugama, M., & Bartholomew, B. (2018). The Arp8 and Arp4 module acts as a DNA sensor controlling INO80 chromatin remodeling. Nat Commun, 9(1), 3309. doi:10.1038/s41467-018-05710-7

Chacin, E., Bansal, P., Reusswig, K. U., Diaz-Santin, L. M., Ortega, P., Vizjak, P., … Kurat, C. F. (2021). A CDK-regulated chromatin segregase promoting chromosome replication. Nat Commun, 12(1), 5224. doi:10.1038/s41467-021-25424-7

Chacin, E., Reusswig, K. U., Furtmeier, J., Bansal, P., Karl, L. A., Pfander, B., … Kurat, C. F. (2023). Establishment and function of chromatin organization at replication origins. Nature, 616(7958), 836–842. doi:10.1038/s41586-023-05926-8

Challa, K., Fajish, V. G., Shinohara, M., Klein, F., Gasser, S. M., & Shinohara, A. (2019). Meiosis-specific prophase-like pathway controls cleavage-independent release of cohesin by Wapl phosphorylation. PLoS Genet, 15(1), e1007851. doi:10.1371/journal.pgen.1007851

Costa, A., & Diffley, J. F. X. (2022). The Initiation of Eukaryotic DNA Replication. Annu Rev Biochem, 91, 107–131. doi:10.1146/annurev-biochem-072321-110228

Creager, R. L., Li, Y., & MacAlpine, D. M. (2015). SnapShot: Origins of DNA replication. Cell, 161(2), 418–418 e411. doi:10.1016/j.cell.2015.03.043

Czajkowsky, D. M., Liu, J., Hamlin, J. L., & Shao, Z. (2008). DNA combing reveals intrinsic temporal disorder in the replication of yeast chromosome VI. J Mol Biol, 375(1), 12–19. doi:10.1016/j.jmb.2007.10.046

Dimitrova, D. S., & Gilbert, D. M. (1999). The spatial position and replication timing of chromosomal domains are both established in early G1 phase. Mol Cell, 4(6), 983–993. doi:10.1016/s1097-2765(00)80227-0

Donaldson, A. D., Fangman, W. L., & Brewer, B. J. (1998). Cdc7 is required throughout the yeast S phase to activate replication origins. Genes Dev, 12(4), 491–501. doi:10.1101/gad.12.4.491

Duncker, B. P., Shimada, K., Tsai-Pflugfelder, M., Pasero, P., & Gasser, S. M. (2002). An N-terminal domain of Dbf4p mediates interaction with both origin recognition complex (ORC) and Rad53p and can deregulate late origin firing. Proc Natl Acad Sci U S A, 99(25), 16087–16092. doi:10.1073/pnas.252093999

Eaton, M. L., Galani, K., Kang, S., Bell, S. P., & MacAlpine, D. M. (2010). Conserved nucleosome positioning defines replication origins. Genes Dev, 24(8), 748–753. doi:10.1101/gad.1913210

Enserink, J. M., & Kolodner, R. D. (2010). An overview of Cdk1-controlled targets and processes. Cell Div, 5, 11. doi:10.1186/1747-1028-5-11

Eustermann, S., Schall, K., Kostrewa, D., Lakomek, K., Strauss, M., Moldt, M., & Hopfner, K. P. (2018). Structural basis for ATP-dependent chromatin remodelling by the INO80 complex. Nature, 556(7701), 386–390. doi:10.1038/s41586-018-0029-y

Fang, D., Lengronne, A., Shi, D., Forey, R., Skrzypczak, M., Ginalski, K., … Lou, H. (2017). Dbf4 recruitment by forkhead transcription factors defines an upstream rate-limiting step in determining origin firing timing. Genes Dev, 31(23-24), 2405–2415. doi:10.1101/gad.306571.117

Fragkos, M., Ganier, O., Coulombe, P., & Mechali, M. (2015). DNA replication origin activation in space and time. Nat Rev Mol Cell Biol, 16(6), 360–374. doi:10.1038/nrm4002

Galanti, L., Peritore, M., Gnugge, R., Cannavo, E., Heipke, J., Palumbieri, M. D., … Pfander, B. (2024). Dbf4-dependent kinase promotes cell cycle controlled resection of DNA double-strand breaks and repair by homologous recombination. Nat Commun, 15(1), 2890. doi:10.1038/s41467-024-46951-z

Gerhold, C. B., Winkler, D. D., Lakomek, K., Seifert, F. U., Fenn, S., Kessler, B., … Hopfner, K. P. (2012). Structure of Actin-related protein 8 and its contribution to nucleosome binding. Nucleic Acids Res, 40(21), 11036–11046. doi:10.1093/nar/gks842

Gillespie, P. J., & Blow, J. J. (2022). DDK: The Outsourced Kinase of Chromosome Maintenance. Biology (Basel), 11(6). doi:10.3390/biology11060877

Greiwe, J. F., Miller, T. C. R., Locke, J., Martino, F., Howell, S., Schreiber, A., … Costa, A. (2022). Structural mechanism for the selective phosphorylation of DNA-loaded MCM double hexamers by the Dbf4-dependent kinase. Nature Structural & Molecular Biology, 29(1), 10–20. doi:10.1038/s41594-021-00698-z

Hawkins, M., Retkute, R., Muller, C. A., Saner, N., Tanaka, T. U., de Moura, A. P., & Nieduszynski, C. A. (2013). High-resolution replication profiles define the stochastic nature of genome replication initiation and termination. Cell Rep, 5(4), 1132–1141. doi:10.1016/j.celrep.2013.10.014

He, W., Rao, H., Tang, S., Bhagwat, N., Kulkarni, D. S., Ma, Y., … Hunter, N. (2020). Regulated Proteolysis of MutSgamma Controls Meiotic Crossing Over. Mol Cell, 78(1), 168–183 e165. doi:10.1016/j.molcel.2020.02.001

Hereford, L. M., & Hartwell, L. H. (1974). Sequential gene function in the initiation of Saccharomyces cerevisiae DNA synthesis. J Mol Biol, 84(3), 445–461. doi:10.1016/0022-2836(74)90451-3

Jiang, W., McDonald, D., Hope, T. J., & Hunter, T. (1999). Mammalian Cdc7-Dbf4 protein kinase complex is essential for initiation of DNA replication. Embo j, 18(20), 5703–5713. doi:10.1093/emboj/18.20.5703

Knoll, K. R., Eustermann, S., Niebauer, V., Oberbeckmann, E., Stoehr, G., Schall, K., … Hopfner, K. P. (2018). The nuclear actin-containing Arp8 module is a linker DNA sensor driving INO80 chromatin remodeling. Nat Struct Mol Biol, 25(9), 823–832. doi:10.1038/s41594-018-0115-8

Kornberg, R. D., & Lorch, Y. (1999). Twenty-five years of the nucleosome, fundamental particle of the eukaryote chromosome. Cell, 98(3), 285–294. doi:10.1016/s0092-8674(00)81958-3

Krietenstein, N., Wal, M., Watanabe, S., Park, B., Peterson, C. L., Pugh, B. F., & Korber, P. (2016). Genomic Nucleosome Organization Reconstituted with Pure Proteins. Cell, 167(3), 709–721 e712. doi:10.1016/j.cell.2016.09.045

Kunert, F., Metzner, F. J., Jung, J., Hopfler, M., Woike, S., Schall, K., … Hopfner, K. P. (2022). Structural mechanism of extranucleosomal DNA readout by the INO80 complex. Sci Adv, 8(49), eadd3189. doi:10.1126/sciadv.add3189

Kurat, C. F., Yeeles, J. T. P., Patel, H., Early, A., & Diffley, J. F. X. (2017). Chromatin Controls DNA Replication Origin Selection, Lagging-Strand Synthesis, and Replication Fork Rates. Mol Cell, 65(1), 117–130. doi:10.1016/j.molcel.2016.11.016

Leonard, A. C., & Mechali, M. (2013). DNA replication origins. Cold Spring Harb Perspect Biol, 5(10), a010116. doi:10.1101/cshperspect.a010116

Liachko, I., Youngblood, R. A., Keich, U., & Dunham, M. J. (2013). High-resolution mapping, characterization, and optimization of autonomously replicating sequences in yeast. Genome Res, 23(4), 698–704. doi:10.1101/gr.144659.112

Lopez-Mosqueda, J., Maas, N. L., Jonsson, Z. O., Defazio-Eli, L. G., Wohlschlegel, J., & Toczyski, D. P. (2010). Damage-induced phosphorylation of Sld3 is important to block late origin firing. Nature, 467(7314), 479–483. doi:10.1038/nature09377

Matos, J., Lipp, J. J., Bogdanova, A., Guillot, S., Okaz, E., Junqueira, M., … Zachariae, W. (2008). Dbf4-dependent CDC7 kinase links DNA replication to the segregation of homologous chromosomes in meiosis I. Cell, 135(4), 662–678. doi:10.1016/j.cell.2008.10.026

Morgan, D. O. (1995). Principles of CDK regulation. Nature, 374(6518), 131–134. doi:10.1038/374131a0

Nieduszynski, C. A., Knox, Y., & Donaldson, A. D. (2006). Genome-wide identification of replication origins in yeast by comparative genomics. Genes Dev, 20(14), 1874–1879. doi:10.1101/gad.385306

Oberbeckmann, E., Niebauer, V., Watanabe, S., Farnung, L., Moldt, M., Schmid, A., … Korber, P. (2021). Ruler elements in chromatin remodelers set nucleosome array spacing and phasing. Nat Commun, 12(1), 3232. doi:10.1038/s41467-021-23015-0

Ocampo, J., Chereji, R. V., Eriksson, P. R., & Clark, D. J. (2019). Contrasting roles of the RSC and ISW1/CHD1 chromatin remodelers in RNA polymerase II elongation and termination. Genome Res, 29(3), 407–417. doi:10.1101/gr.242032.118

On, K. F., Beuron, F., Frith, D., Snijders, A. P., Morris, E. P., & Diffley, J. F. (2014). Prereplicative complexes assembled in vitro support origin-dependent and independent DNA replication. Embo j, 33(6), 605–620. doi:10.1002/embj.201387369

Papamichos-Chronakis, M., & Peterson, C. L. (2008). The Ino80 chromatin-remodeling enzyme regulates replisome function and stability. Nat Struct Mol Biol, 15(4), 338–345. doi:10.1038/nsmb.1413

Pasero, P., Duncker, B. P., Schwob, E., & Gasser, S. M. (1999). A role for the Cdc7 kinase regulatory subunit Dbf4p in the formation of initiation-competent origins of replication. Genes Dev, 13(16), 2159–2176. doi:10.1101/gad.13.16.2159

Princz, L. N., Wild, P., Bittmann, J., Aguado, F. J., Blanco, M. G., Matos, J., & Pfander, B. (2017). Dbf4-dependent kinase and the Rtt107 scaffold promote Mus81-Mms4 resolvase activation during mitosis. Embo j, 36(5), 664–678. doi:10.15252/embj.201694831

Raghuraman, M. K., Brewer, B. J., & Fangman, W. L. (1997). Cell cycle-dependent establishment of a late replication program. Science, 276(5313), 806–809. doi:10.1126/science.276.5313.806

Reusswig, K. U., Zimmermann, F., Galanti, L., & Pfander, B. (2016). Robust Replication Control Is Generated by Temporal Gaps between Licensing and Firing Phases and Depends on Degradation of Firing Factor Sld2. Cell Rep, 17(2), 556–569. doi:10.1016/j.celrep.2016.09.013

Rossi, M. J., Kuntala, P. K., Lai, W. K. M., Yamada, N., Badjatia, N., Mittal, C., … Pugh, B. F. (2021). A high-resolution protein architecture of the budding yeast genome. Nature, 592(7853), 309–314. doi:10.1038/s41586-021-03314-8

Safaric, B., Chacin, E., Scherr, M. J., Rajappa, L., Gebhardt, C., Kurat, C. F., … Duderstadt, K. E. (2022). The fork protection complex recruits FACT to reorganize nucleosomes during replication. Nucleic Acids Res, 50(3), 1317–1334. doi:10.1093/nar/gkac005

Saner, N., Karschau, J., Natsume, T., Gierlinski, M., Retkute, R., Hawkins, M., … Tanaka, T. U. (2013). Stochastic association of neighboring replicons creates replication factories in budding yeast. J Cell Biol, 202(7), 1001–1012. doi:10.1083/jcb.201306143

Saravanan, M., Wuerges, J., Bose, D., McCormack, E. A., Cook, N. J., Zhang, X., & Wigley, D. B. (2012). Interactions between the nucleosome histone core and Arp8 in the INO80 chromatin remodeling complex. Proc Natl Acad Sci U S A, 109(51), 20883–20888. doi:10.1073/pnas.1214735109

Sasanuma, H., Hirota, K., Fukuda, T., Kakusho, N., Kugou, K., Kawasaki, Y., … Ohta, K. (2008). Cdc7-dependent phosphorylation of Mer2 facilitates initiation of yeast meiotic recombination. Genes Dev, 22(3), 398–410. doi:10.1101/gad.1626608

Sheu, Y. J., & Stillman, B. (2006). Cdc7-Dbf4 phosphorylates MCM proteins via a docking site-mediated mechanism to promote S phase progression. Mol Cell, 24(1), 101–113. doi:10.1016/j.molcel.2006.07.033

Sheu, Y. J., & Stillman, B. (2010). The Dbf4-Cdc7 kinase promotes S phase by alleviating an inhibitory activity in Mcm4. Nature, 463(7277), 113–117. doi:10.1038/nature08647

Singh, A. K., Schauer, T., Pfaller, L., Straub, T., & Mueller-Planitz, F. (2021). The biogenesis and function of nucleosome arrays. Nature Communications, 12(1), 7011. doi:10.1038/s41467-021-27285-6

Siow, C. C., Nieduszynska, S. R., Muller, C. A., & Nieduszynski, C. A. (2012). OriDB, the DNA replication origin database updated and extended. Nucleic Acids Res, 40(Database issue), D682-686. doi:10.1093/nar/gkr1091

Smith, D. J., & Whitehouse, I. (2012). Intrinsic coupling of lagging-strand synthesis to chromatin assembly. Nature, 483(7390), 434–438. doi:10.1038/nature10895

Soriano, I., Morafraile, E. C., Vazquez, E., Antequera, F., & Segurado, M. (2014). Different nucleosomal architectures at early and late replicating origins in Saccharomyces cerevisiae. BMC Genomics, 15(1), 791. doi:10.1186/1471-2164-15-791

Szerlong, H., Hinata, K., Viswanathan, R., Erdjument-Bromage, H., Tempst, P., & Cairns, B. R. (2008). The HSA domain binds nuclear actin-related proteins to regulate chromatin-remodeling ATPases. Nat Struct Mol Biol, 15(5), 469–476. doi:10.1038/nsmb.1403

Szklarczyk, D., Kirsch, R., Koutrouli, M., Nastou, K., Mehryary, F., Hachilif, R., … von Mering, C. (2023). The STRING database in 2023: protein-protein association networks and functional enrichment analyses for any sequenced genome of interest. Nucleic Acids Res, 51(D1), D638–d646. doi:10.1093/nar/gkac1000

Tkach, J. M., Yimit, A., Lee, A. Y., Riffle, M., Costanzo, M., Jaschob, D., … Brown, G. W. (2012). Dissecting DNA damage response pathways by analysing protein localization and abundance changes during DNA replication stress. Nature Cell Biology, 14(9), 966–976. doi:10.1038/ncb2549

Tosi, A., Haas, C., Herzog, F., Gilmozzi, A., Berninghausen, O., Ungewickell, C., … Hopfner, K. P. (2013). Structure and subunit topology of the INO80 chromatin remodeler and its nucleosome complex. Cell, 154(6), 1207–1219. doi:10.1016/j.cell.2013.08.016

Ubersax, J. A., Woodbury, E. L., Quang, P. N., Paraz, M., Blethrow, J. D., Shah, K., … Morgan, D. O. (2003). Targets of the cyclin-dependent kinase Cdk1. Nature, 425(6960), 859–864. doi:10.1038/nature02062

Vincent, J. A., Kwong, T. J., & Tsukiyama, T. (2008). ATP-dependent chromatin remodeling shapes the DNA replication landscape. Nat Struct Mol Biol, 15(5), 477–484. doi:10.1038/nsmb.1419

Wan, L., Niu, H., Futcher, B., Zhang, C., Shokat, K. M., Boulton, S. J., & Hollingsworth, N. M. (2008). Cdc28-Clb5 (CDK-S) and Cdc7-Dbf4 (DDK) collaborate to initiate meiotic recombination in yeast. Genes Dev, 22(3), 386–397. doi:10.1101/gad.1626408

Xu, W., Aparicio, J. G., Aparicio, O. M., & Tavare, S. (2006). Genome-wide mapping of ORC and Mcm2p binding sites on tiling arrays and identification of essential ARS consensus sequences in S. cerevisiae. BMC Genomics, 7, 276. doi:10.1186/1471-2164-7-276

Zegerman, P., & Diffley, J. F. (2010). Checkpoint-dependent inhibition of DNA replication initiation by Sld3 and Dbf4 phosphorylation. Nature, 467(7314), 474–478. doi:10.1038/nature09373

Zhou, C. Y., Johnson, S. L., Gamarra, N. I., & Narlikar, G. J. (2016). Mechanisms of ATP-Dependent Chromatin Remodeling Motors. Annu Rev Biophys, 45, 153–181. doi:10.1146/annurev-biophys-051013-022819

Zhou, C. Y., Johnson, S. L., Lee, L. J., Longhurst, A. D., Beckwith, S. L., Johnson, M. J., … Narlikar, G. J. (2018). The Yeast INO80 Complex Operates as a Tunable DNA Length-Sensitive Switch to Regulate Nucleosome Sliding. Mol Cell, 69(4), 677–688 e679. doi:10.1016/j.molcel.2018.01.028

## Methods References

Aftab, W., Lahiri, S., & Imhof, A. (2022). ImShot: An Open-Source Software for Probabilistic Identification of Proteins In Situ and Visualization of Proteomics Data. Mol Cell Proteomics, 21(6), 100242. doi:10.1016/j.mcpro.2022.100242

Almer, A., & Hörz, W. (1986). Nuclease hypersensitive regions with adjacent positioned nucleosomes mark the gene boundaries of the PHO5/PHO3 locus in yeast. Embo j, 5(10), 2681–2687. doi:10.1002/j.1460-2075.1986.tb04551.x

Barrios, A., Selleck, W., Hnatkovich, B., Kramer, R., Sermwittayawong, D., & Tan, S. (2007). Expression and purification of recombinant yeast Ada2/Ada3/Gcn5 and Piccolo NuA4 histone acetyltransferase complexes. Methods, 41(3), 271–277. doi:10.1016/j.ymeth.2006.08.007

Biswas, D., Yu, Y., Prall, M., Formosa, T., & Stillman, D. J. (2005). The yeast FACT complex has a role in transcriptional initiation. Mol Cell Biol, 25(14), 5812–5822. doi:10.1128/mcb.25.14.5812-5822.2005

Brachmann, C. B., Davies, A., Cost, G. J., Caputo, E., Li, J., Hieter, P., & Boeke, J. D. (1998). Designer deletion strains derived from Saccharomyces cerevisiae S288C: a useful set of strains and plasmids for PCR-mediated gene disruption and other applications. Yeast, 14(2), 115–132. doi:10.1002/(sici)1097-0061(19980130)14:2<115::Aid-yea204>3.0.Co;2-2

Combe, C. W., Fischer, L., & Rappsilber, J. (2015). xiNET: cross-link network maps with residue resolution. Mol Cell Proteomics, 14(4), 1137–1147. doi:10.1074/mcp.O114.042259

Dyer, P. N., Edayathumangalam, R. S., White, C. L., Bao, Y., Chakravarthy, S., Muthurajan, U. M., & Luger, K. (2003). Reconstitution of Nucleosome Core Particles from Recombinant Histones and DNA. In Methods in Enzymology (Vol. 375, pp. 23–44): Academic Press.

Gómez-González, B., Ruiz, J. F., & Aguilera, A. (2011). Genetic and molecular analysis of mitotic recombination in Saccharomyces cerevisiae. Methods Mol Biol, 745, 151–172. doi:10.1007/978-1-61779-129-1_10

Humphrey, S. J., Karayel, O., James, D. E., & Mann, M. (2018). High-throughput and high-sensitivity phosphoproteomics with the EasyPhos platform. Nat Protoc, 13(9), 1897–1916. doi:10.1038/s41596-018-0014-9

Ihling, C. H., Piersimoni, L., Kipping, M., & Sinz, A. (2021). Cross-Linking/Mass Spectrometry Combined with Ion Mobility on a timsTOF Pro Instrument for Structural Proteomics. Anal Chem, 93(33), 11442–11450. doi:10.1021/acs.analchem.1c01317

Kingston, I. J., Yung, J. S., & Singleton, M. R. (2011). Biophysical characterization of the centromere-specific nucleosome from budding yeast. J Biol Chem, 286(5), 4021–4026. doi:10.1074/jbc.M110.189340

Ludwigsen, J., Hepp, N., Klinker, H., Pfennig, S., & Mueller-Planitz, F. (2018). Remodeling and Repositioning of Nucleosomes in Nucleosomal Arrays. Methods Mol Biol, 1805, 349–370. doi:10.1007/978-1-4939-8556-2_18

Rappsilber, J., Mann, M., & Ishihama, Y. (2007). Protocol for micro-purification, enrichment, pre-fractionation and storage of peptides for proteomics using StageTips. Nat Protoc, 2(8), 1896–1906. doi:10.1038/nprot.2007.261

Reusswig, K. U., Bittmann, J., Peritore, M., Courtes, M., Pardo, B., Wierer, M., … Pfander, B. (2022). Unscheduled DNA replication in G1 causes genome instability and damage signatures indicative of replication collisions. Nat Commun, 13(1), 7014. doi:10.1038/s41467-022-34379-2

Shen, X. (2004). Preparation and analysis of the INO80 complex. Methods Enzymol, 377, 401–412. doi:10.1016/s0076-6879(03)77026-8

Winston, F., Dollard, C., & Ricupero-Hovasse, S. L. (1995). Construction of a set of convenient Saccharomyces cerevisiae strains that are isogenic to S288C. Yeast, 11(1), 53–55. doi:10.1002/yea.320110107

Yılmaz, Ş., Busch, F., Nagaraj, N., & Cox, J. (2022). Accurate and Automated High-Coverage Identification of Chemically Cross-Linked Peptides with MaxLynx. Anal Chem, 94(3), 1608–1617. doi:10.1021/acs.analchem.1c03688

Zhang, X., Smits, A. H., van Tilburg, G. B. A., Ovaa, H., Huber, W., & Vermeulen, M. (2018). Proteome-wide identification of ubiquitin interactions using UbIA-MS. Nature Protocols, 13(3), 530–550. doi:10.1038/nprot.2017.147

